# A circuit of protein-protein regulatory interactions enables polarity establishment in a bacterium

**DOI:** 10.1101/503250

**Authors:** Wei Zhao, Samuel W. Duvall, Kimberly A. Kowallis, Dylan T. Tomares, Haley N. Petitjean, W. Seth Childers

**Author notes:** Lead Contact: W. Seth Childers.

## Abstract

Asymmetric cell division generates specialized daughter cells that play a variety of roles including tissue morphogenesis in eukaryotes and pathogenesis in bacteria. In the gram-negative bacterium *Caulobacter crescentus*, asymmetric localization of two biochemically distinct signaling hubs at opposite cell poles provides the foundation for asymmetric cell division. Through a set of genetic, synthetic biology and biochemical approaches we have characterized the regulatory interactions between three scaffolding proteins. These studies have revealed that the scaffold protein PodJ functions as a central mediator for organizing the new cell signaling hub, including promoting bipolarization of the central developmental scaffold protein PopZ. In addition, we identified that the old pole scaffold SpmX serves as a negative regulator of PodJ subcellular accumulation. These two scaffold-scaffold regulatory interactions serve as the core of an integrated cell polarization circuit that is layered on top of the cell-cycle circuitry to coordinate cell differentiation and asymmetric cell division.

## Introduction

The earliest stage of an asymmetric cell division is the unequal inheritance of cell fate determinants. While models of eukaryotic cell polarity have been developed (Chau et al., 2012) it remains unclear what mechanisms are employed by bacteria to achieve polarity (McAdams and Shapiro, 2009). The degree of asymmetry of cell division in the bacterial kingdom has only been sparsely examined, however broad implications in pathogenesis (Van der Henst et al., 2013) and persister cell development (Aakre and Laub, 2012) have been reported.

*Caulobacter crescentus* is a well-established bacterial model organism for examining how bacterial cells divide asymmetrically (Shapiro et al., 1971). The cell division results in two morphologically and functionally distinct cells (Figure 1A): a motile swarmer cell that is incapable of chromosome replication and a sessile stalked cell that initiates replication once per cell-cycle (Bergé and Viollier, 2018; Curtis and Brun, 2010; Lasker et al., 2016). These two cell-types utilize bimodal survival strategies, including distinct responses to heavy metal stress (Lawaree et al., 2016) and differences in buoyancy (Ardissone et al., 2014). The swarmer cell can differentiate into stalked cell by shedding the flagellum and initiating stalk biogenesis and chromosome replication (Figure 1A). Genetic studies have revealed that *C. crescentus* bacteria do not use the primary cell polarity regulators that drive eukaryotic stem cell division, but use a set of self-assembled scaffolding proteins to achieve these goals (Bergé and Viollier, 2018; Lasker et al., 2016; Tsokos and Laub, 2012).

**Figure 1:**
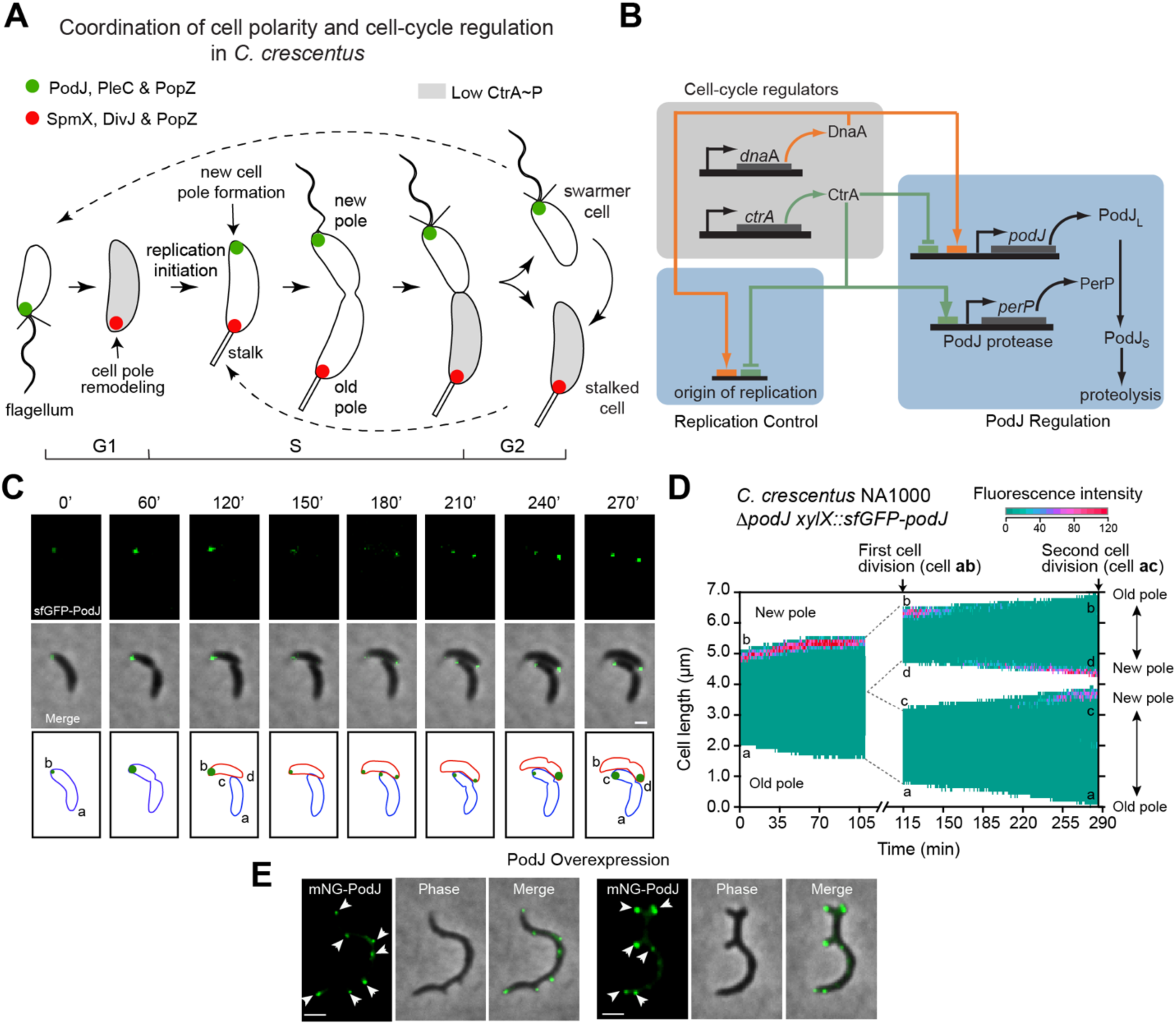
The scaffolding protein PodJ recognizes the new pole and is involved in cell polarity establishment. (A) Schematic of cell polarity establishment and cell cycle regulation in *Caulobacter crescentus*. Swarmer cells differentiate into stalked cells, which is correlated with cell pole remodeling of a PodJ-rich signaling hub (green) into a SpmX-rich signaling hub (red). In stalked cells, after initiation of replication a PodJ-rich signaling hub accumulates at the new cell pole. Cell division results in a swarmer cell that involved unequal inheritance of a PodJ-rich signaling hub in swarmer cells and a SpmX-rich signaling hub in stalked cells. (B) Two key cell-cycle regulators, CtrA and DnaA, co-regulate the initiation of replication and the transcription of PodJ and a PodJ specific protease PerP (C and D) Time-lapse microscopy analyses show PodJ accumulates at the new cell pole during the cell cycle. Kymographs of sfGFP-PodJ signal along the cell length over time after the synchronization of WSC1201 swarmer cells, images were acquired every 2 min. (E) Constitutive overnight PodJ overexpression (0.3% xylose) causes formation of ectopic cell poles that are co-localized with PodJ foci. Two representative cells are shown. All bars, 2 μm.

Much of the developmental differences between the *C. crescentus* swarmer and stalked cell-types is coordinated by the CtrA signaling pathway which regulates more than 90 genes associated with flagella and stalk biogenesis, chromosome replication and cell wall growth (Laub et al., 2002). Asymmetric activation of the CtrA signaling pathway arises due to the asymmetric localization of two compositionally and functionally distinct signaling complexes (Lasker et al., 2016; Matroule et al., 2004). In pre-divisional cells the new cell pole signaling hub is composed of three scaffolding proteins (TipN (Huitema et al., 2006; Lam et al., 2006), PodJ (Hinz et al., 2003; Viollier et al., 2002) and PopZ (Bowman et al., 2008; Ebersbach et al., 2008)) and eight key signaling proteins (PleC, CpaE, DivL, DivK, PleD, PopA, MopJ and CckA) that function together to activate the CtrA pathway via phosphorylation (Lasker et al., 2016). The old cell pole is composed of two scaffolds (SpmX (Radhakrishnan et al., 2008) and PopZ) and three signaling proteins (DivJ, DivK and PleD) that work as a concerted system to deactivate CtrA through both dephosphorylation and proteolysis (Lasker et al., 2016). The positioning of these complexes prior to division ensures that the daughter cells differentially regulate CtrA and develop as unique cell-types (Figure 1A). Moreover, as newborn swarmer cells differentiate into stalked cells, their inherited PodJ-rich signaling hub undergoes compositional remodeling to become a SpmX-rich signaling hub (Figure 1A). Central questions remain regarding how *C. crescentus* cells establish, maintain and remodel cell polarity to coordinate asymmetric cell division with the cell-cycle.

The PopZ scaffolding protein plays a central role in organizing the two organelle-like signaling complexes, as it binds directly to seven client proteins (Holmes et al., 2016). However, PopZ is a common scaffold shared by both the new and old cell pole signaling hubs. This raises the question of how PopZ prevents mixing of signaling hubs when it switches from a monopolar to a bipolar localization pattern as cell develop from a swarmer cell into a pre-divisional cell (Figure 1A). Studies have shown that timing of new cell pole accumulation of PopZ is correlated with the initiation of replication (Laloux and Jacobs-Wagner, 2013; Mera et al., 2014). Studies have indicated that the coordination of cell cycle and cell polarity could be mediated by proteins whose transcription is activated by DnaA and repressed by CtrA (Figure 1B) (Crymes et al., 1999). Two proteins contribute to PopZ bipolarization, ZitP (Berge et al., 2016) and TipN (Laloux and Jacobs-Wagner, 2013), however each of these studies suggest that redundant factors promote PopZ’s new cell pole binding. One potential candidate protein is the new cell pole signaling scaffold PodJ, whose transcription upregulated at the same time as the initiation of replication (Figure 1B) (Crymes et al., 1999) and is proteolyzed prior to the swarmer-to-stalk transition (Chen et al., 2006; Curtis et al., 2012). Meanwhile, PodJ is required for establishment of cell polarity and asymmetric cell division (Hinz et al., 2003). PodJ directly or indirectly recruits the PleC kinase (Curtis et al., 2012; Hinz et al., 2003; Viollier et al., 2002), the pilus assembly protein CpaE (Viollier et al., 2002), and the ClpXP protease adaptor protein PopA (Duerig et al., 2009; Ozaki et al., 2014) to the new cell pole.

The PodJ scaffold domain architecture includes a N-terminal cytosolic domain composed of a coiled-coil rich region (Lawler et al., 2006) that is adjacent to an intrinsically disordered region (Figure 2A). The disordered region is comprised of two compositionally distinct sections. Residues 471-588 are rich in proline, serine and glutamic acid residues with a net charge of −18. While residues 589-642 are rich in serine, glycine and lysine residues with a net charge of +10. The C-terminal end passes through the membrane into the periplasm and contains a tetrapeptide co-repeat domain and a peptidoglycan binding domain (Lawler et al., 2006). The periplasmic domains coordinate pili biogenesis at the new cell pole and have been shown to be dispensable for PodJ localization at the cell pole (Lawler et al., 2006). PodJ’s cytoplasmic domains have been implicated directly or indirectly with the recruitment of signaling proteins that activate the CtrA pathway (PleC (Curtis et al., 2012; Lawler et al., 2006; Viollier et al., 2002), DivL (Curtis et al., 2012)), initiate pili biogenesis (CpaE (Curtis et al., 2012; Viollier et al., 2002)) and localize the holdfast complex (HfaB (Hardy et al., 2010)). Previous domain analysis has implicated a portion of the intrinsically disordered domain and the periplasmic domain in new cell pole targeting of PodJ (Lawler et al., 2006). However it remains unclear if this cell pole recruitment is dependent upon other polarity proteins (e.g. TipN or PopZ), and more broadly how the PodJ contributes to polarity establishment. Here we apply a combination of heterologous reconstitution experiments, genetics, quantitative cell biology and biochemical assays to map the regulatory interactions between three central scaffolding proteins (PopZ, PodJ and SpmX) that can account for exquisite cell polarity observed in asymmetrically dividing *C. crescentus* cells.

**Figure 2:**
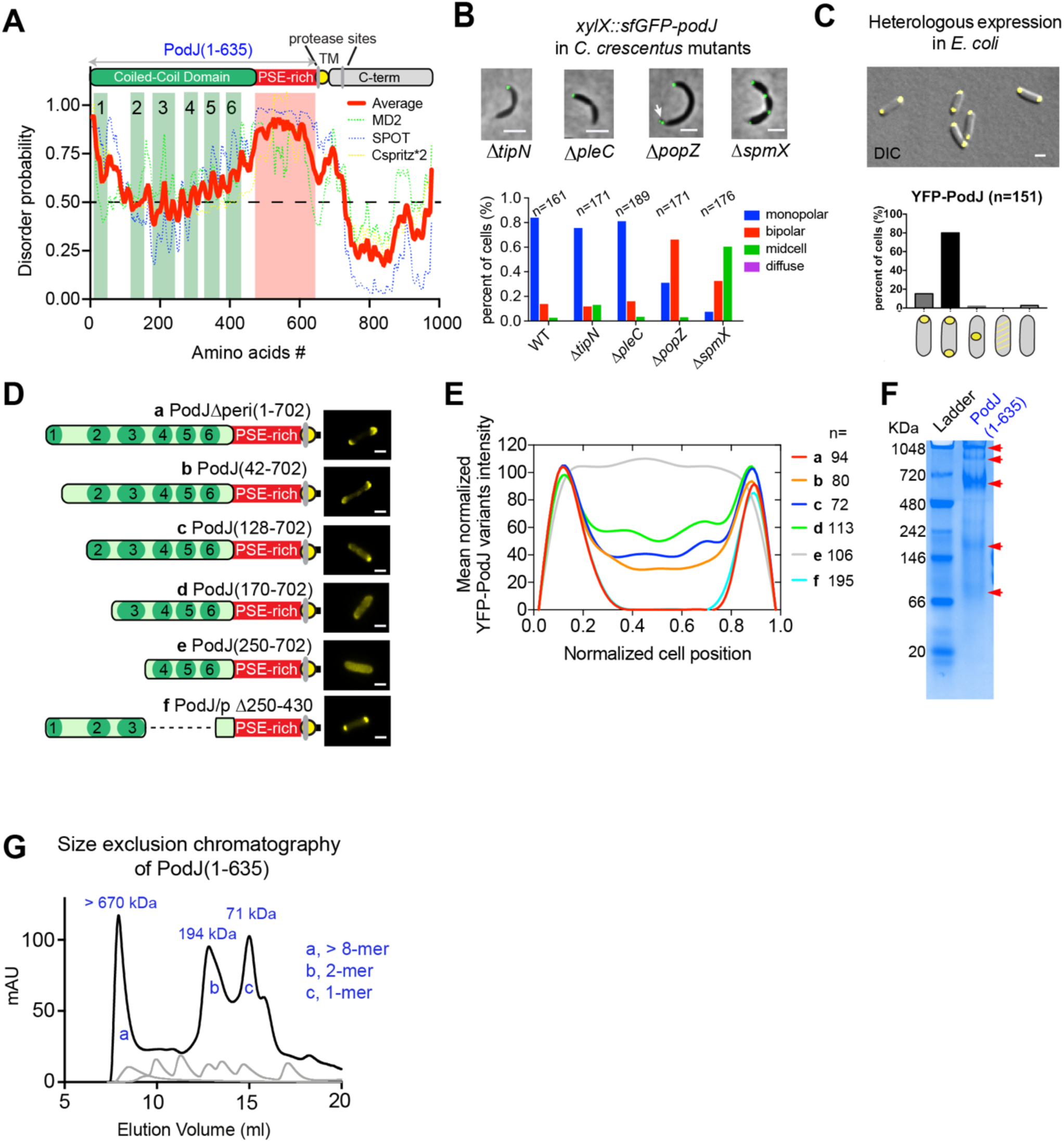
PodJ is a self-assembled protein, whose monopolar accumulation is dependent upon PodJ and SpmX. (A) PodJ domain organization predicted by HHpred and adapted from previous studies(Curtis et al., 2012; Lawler et al., 2006) The coiled-coil rich region was analyzed by PCOILS and modeled with MODELLER. The probability of intrinsic disorder over the primary sequence of PodJ (red line), represented as the average scores from four disorder prediction algorithms: Meta disorder MD2, SPOT, Cspritz*2, and MFDp2. (B) PodJ accumulation as a concentrated focus in *C. crescentus* cells is independent of TipN, PleC, PopZ and SpmX. However, PodJ monopolar subcellular accumulation is dependent upon the PopZ and SpmX scaffolds. (C) Subcellular localization of PodJ when heterologously expressed YFP-PodJ in *E. coli*. (D) Heterologous expression of YFP-PodJ variants in *E. coli* reveals that the N-terminal coiled-coil region 1-3 is critical for accumulation of PodJ at the cell poles. (E) The signal of YFP-PodJ along the cell length was plotted for the cells shown in Figure 2D. All of the data was normalized with the highest intensity in each strain setting as 100%. (F) Purified PodJ(1-635) was analyzed *in vitro* via native gel analysis. The protein is subjected to nondenaturing gel electrophoresis at 4°C and subsequently stained with Coomassie blue stain. Four distinct bands indicate PodJ oligomers larger 480 kDa. (G) Analytical size exclusion chromatography was used to measure the apparent molecular mass of purified PodJ(1-635) (black line), by plotting absorbance at 215 nm versus elution volume. The indicated molecular masses of each peak were determined by comparison to the elution volume of protein standards (grey lines). A representative trace is shown from three independent replicates.

## Results

### PodJ expression level is critical for the maintenance of cell polarity

Spatiotemporal PodJ protein abundance is highly regulated at the levels of transcription (Crymes et al., 1999) and proteolysis (Chen et al., 2006; Chen et al., 2005; Curtis et al., 2012) (Figure 1B). Expression of sfGFP-PodJ at a 0.03% xylose induction level in a Δ*podJ* strain resulted in PodJ accumulation at the new cell pole (Figure 1C and 1D). During the swarmer-to-stalked cell transition, the PodJ focus at the old cell pole diminished, while a new PodJ focus accumulated at the new cell pole (Figure 1C, Figure S1A, Movie S1). This localization pattern is consistent with previous immunofluorescence microscopy observations (Hinz et al., 2003; Viollier et al., 2002). In contrast, overexpression of PodJ resulted in cell filamentation and several small ectopic cell poles that were observed in 53% of cells (Figure 1E), each containing a PodJ focus (Figure 1E). These data indicate that strict regulation of PodJ protein expression levels in *C. crescentus* is critical for maintenance of cell polarity and rod morphology.

### PodJ monopolar subcellular accumulation is dependent upon the old cell pole complex

Live cell imaging of sfGFP-PodJ in Δ*popZ*, Δ*spmX*, Δ*tipN* or Δ*pleC* strains showed that PodJ accumulated at the cell poles in each of these deletion strain (Figure 2B). Therefore, PodJ accumulation as a focus was independent of other known scaffold proteins. However, we did observe that PodJ subcellular position was dependent upon PopZ or SpmX, as the percentage of cells containing monopolar PodJ significantly reduced from 82% in wild-type cell to 35% for Δ*popZ*, and 9% for Δ*spmX* (Figure 2B). Because deletion of *spmX* results in long-chain cell phenotype in *C. crescentus* (Radhakrishnan et al., 2008), the accumulated PodJ foci also occupied each constriction sites. Collectively, these results suggest that PodJ accumulation as a focus is independent of other known scaffold proteins. However, the subcellular positioning of PodJ is dependent directly or indirectly upon the scaffolding proteins PopZ and SpmX.

### PodJ accumulates at the cell poles in E. coli independent of specific C. crescentus proteins

To further test if PodJ subcellular accumulation was independent of other *C. crescentus* specific factors, we heterologously expressed PodJ in *Escherichia coli* BL21 cells. Notably, the γ-proteobacterium *E. coli* is highly divergent from the α-proteobacterium *C. crescentus* and does not contain any clear homologs of the *C. crescentus* scaffolding proteins or new cell pole signaling proteins. YFP-PodJ accumulated at both cell poles in *E. coli* (Figure 2C), suggesting that PodJ cell pole accumulation was independent of known *C. crescentus* polarity proteins. Co-expression of YFP-PodJ together with inclusion body marker IbpA-mCherry (Lindner et al., 2008) demonstrated that PodJ did not co-localize with inclusion bodies in *E. coli* (Figure S3B). Moreover, YFP-PodJ localized as a bipolar pattern in about 80% of *E. coli* cells (Figure 2C), which differs from its monopolar localization pattern in *C. crescentus*. These results suggest that possible negative or positive regulators may promote the monopolar accumulation of PodJ in *C. crescentus*.

To identify a minimal cell pole accumulating domain, we screened a set of 21 PodJ domain deletion variants for their capabilities to maintain cell pole accumulation (Figure 2D, Figure S2). Amongst this set, the construct representing the PerP-cleaved form of PodJ that lacks the periplasmic domains, PodJΔperi, accumulated at the cell poles similar to the wild-type PodJ in *E. coli* (Figure 2D), consistent with earlier studies (Lawler et al., 2006). Deletion of the intrinsically disordered PSE-rich region or CC4-6 does not affect PodJ subcellular accumulation (Figure S3). In contrast, the PodJ localization pattern gradually changed from bipolar to diffuse when we truncated the coiled-coil (CC) domains 1 to 3 (Figure 2D), indicating that residues 1-249 were critical for cell pole accumulation of PodJ.

### PodJ self-assembled into a high order oligomer in vitro

Since PodJ accumulated at the cell poles independent of other known scaffolding proteins *in vivo*, we hypothesized that PodJ is a self-assembled protein. We therefore purified the cytoplasmic portion of PodJ, PodJ(1-635), and analyzed the protein through native gel analysis (Figure 2F). The result showed PodJ oligomerization into an array of large oligomeric complexes ranging in size from 720-1048 kDa. Further analytical size exclusion chromatography of PodJ(1-635) indicated PodJ oligomers ranging in size from: 71 kDa (monomer), 194 kDa (dimer), and > 650 kDa (> 8-mer) (Figure 2G). These results indicate that PodJ is a self-assembled scaffolding proteins, and that subcellular localization may be dependent upon coiled-coil multivalent interactions.

### PodJ is the central organizer of new cell pole assembly

Previous studies have suggested that PodJ could serve as a scaffolding protein as PleC (Viollier et al., 2002), CpaE (Ozaki et al., 2014; Viollier et al., 2002), and PopA (Ozaki et al., 2014)) all require PodJ for subcellular accumulation. However, it remains elusive if these dependencies are due to a direct or indirect recruitment by PodJ. To address these questions, we screened 17 cell-cycle regulatory proteins (Figure S2A) for their capacity to co-localize with PodJ upon heterologous co-expression in *E. coli*. Three predicted PodJ interaction partners (PleC, CpaE, and PopA) exhibited mostly diffused pattern when expressed alone, but co-localized with PodJ at the cell poles when co-expressed (Figure 3A, 3B). Three other proteins (SpmX, FtsZ, and TipN) disrupted the PodJ localization pattern when co-expressed with them in *E. coli* (Figure S3C), suggesting they could serve as a negative regulator of PodJ localization. We were unable to detect any direct interactions between PodJ and the following new cell pole signaling proteins: DivL, DivK, CckA, ParA, ParB, and PleD (Figure S3D, Figure 3B). These results support a model of PodJ as a scaffold protein that directly recruits at least three client proteins: PleC, CpaE and PopA.

**Figure 3:**
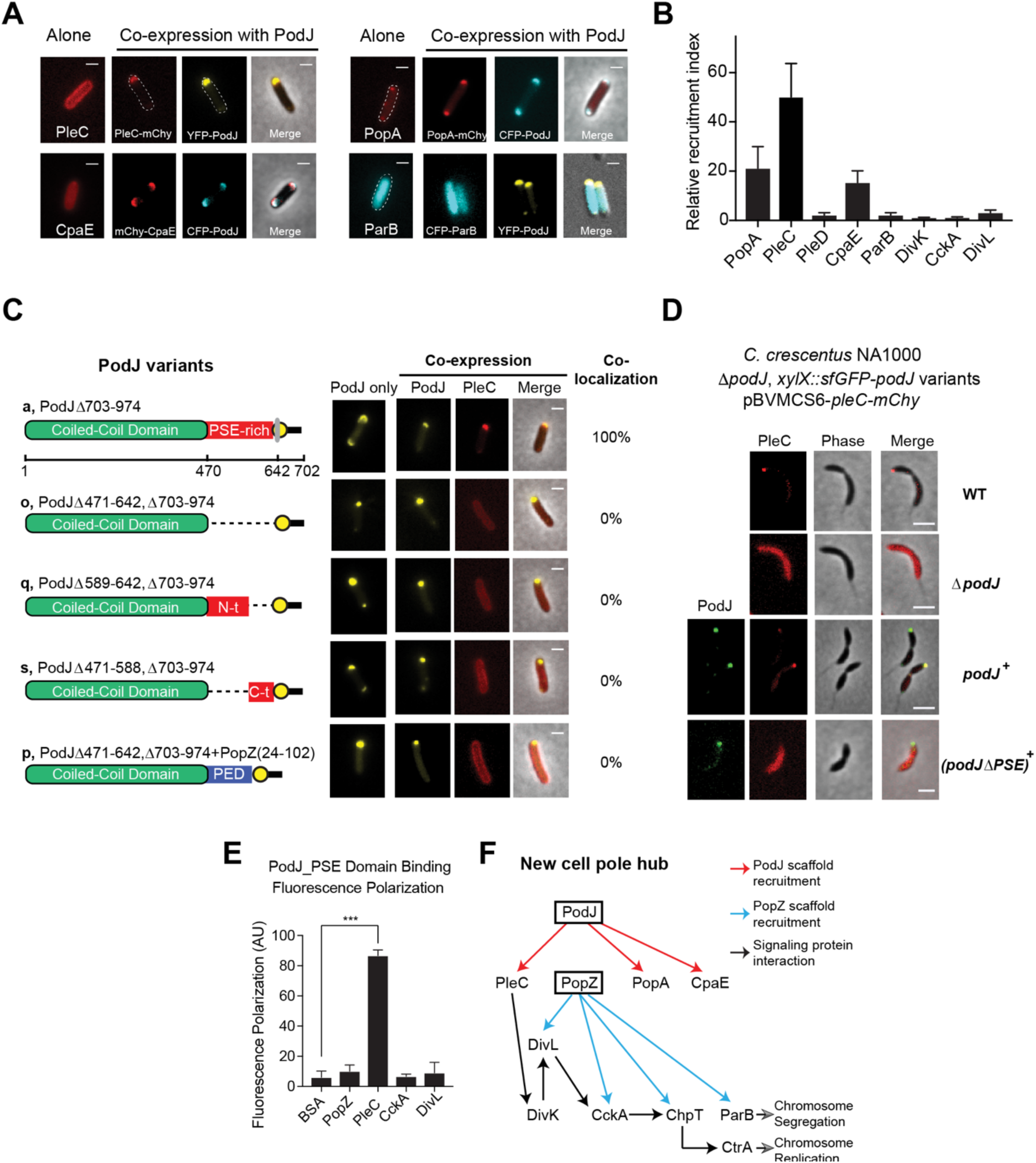
PodJ is a central organizer of the new cell pole signaling hub. The interactions between PodJ and binding partners were evaluated through heterologous co-expression in *E. coli*. (A) YFP-PodJ exhibits selective recruitment roles for the *C. crescentus* new cell pole proteins that include PleC, PopA and CpaE. (B) The polar recruitment ability by PodJ was calculated and normalized as a relative localization index. (C) Co-expression of PodJ variants together with PleC reveals that the intrinsically disordered PSE-rich domain is necessary for PleC recruitment to the cell pole in *E. coli*. (D) Expression of PodJ variants as a sole-copy in *C. crescentus* revealed that the PSE-rich domain is required for PleC accumulation at the new cell poles. (E) Fluorescence polarization binding assay confirms PodJ directly interaction with PleC. 100 nM BODIPY dye labeled PodJ_PSE mixed with the following new cell pole proteins at 10 µM PopZ, PleC, CckA and DivL. PodJ_PSE specifically binds to PleC. (G) New cell pole localization hierarchy based upon these studies indicate that PodJ is a central organizer in addition to PopZ(Holmes et al., 2016) for the new cell pole signaling hub. Interactions identified in this study (red), and those discovered previously by Holmes et al. (blue)(Holmes et al., 2016).

Amongst these proteins, the positioning of PleC at the new cell pole is critical for activation of the CtrA signaling pathway and generation of a CtrA signaling gradient (Chen et al., 2011; Matroule et al., 2004). Two regions of PodJ which contribute additively to the localization of PleC: the C-terminal peptidoglycan binding domain (residues 921-974) and residues 589-639 (Curtis et al., 2012; Lawler et al., 2006). Through the *E. coli* heterologous co-expression assay (Figure S4) we observed that PleC could co-localize robustly with a PodJ variant lacking the C-terminal periplasmic region (PodJΔ703-974, Figure 3C). Deletion of either the N-terminal (PodJΔ471-588) or the C-terminal (PodJ Δ589-642) disordered region resulted in a loss of PleC co-localization in *E. coli* (Figure 3C). These results suggest that the entire disordered region of PodJ serves as a binding site for PleC and may be involved in weak multivalent interactions with this domain. Moreover, we demonstrated that *C. crescentus* cells that express PodJΔPSE as a sole copy display a diffuse PleC subcellular localization pattern in a manner similar to the Δ*podJ* strain (Figure 3D). We next tested if the negatively charged disordered region could be substituted with other disordered proteins, by replacing PodJ’s PSE domain with PopZ’s disordered PED-rich domain. A similar diffuse pattern of PleC was observed in *E. coli* when we swapped the PodJ PSE-rich disordered region with the disordered PED-rich region from PopZ(Holmes et al., 2016) (Figure 3C). Therefore, our results emphasized the PSE domain likely presents a specific site for recruitment of PleC client proteins.

We fluorescently labeled PodJ_PSE domain with a BODIPY dye and measured binding via a fluorescence polarization assay by mixing 16 µM of PleC sensory domain with 100 nM of BODIPY-PodJ_PSE. As shown in Figure 3E, PodJ_PSE bound selectively to PleC, however did not bind to other signaling proteins (e.g. CckA and DivL) that co-localize at the new cell pole. In summary, by integrating our studies together with previous work from PopZ (Holmes et al., 2016), DivL(Mann and Shapiro, 2018; Tsokos et al., 2011), and CckA (Biondi et al., 2006), we have mapped out a localization dependency hierarchy that implicates two scaffolds, PodJ and PopZ, as nucleating factors for new cell pole assembly (Figure 3F).

### PodJ nucleates PopZ assembly at the new cell pole in C. crescentus

PopZ plays a critical role in the predivisional cell polarity establishment by switching from a monopolar to bipolar localization (Bowman et al., 2010; Ptacin et al., 2014) and recruiting core CtrA regulatory proteins: DivL, CckA, and ChpT (Holmes et al., 2016). We tested if PopZ accumulation at the new cell pole was PodJ dependent in *C. crescentus*. We observed that in the Δ*podJ* strain, there was a 4-fold reduction in the fraction of mCherry-PopZ signal at the new cell pole versus wild-type strains (Figure 4A, 4B). Notably, robust PopZ accumulation at the new cell pole could be rescued when sfGFP-*podJ* was expressed from the chromosomal xylose locus (Figure 4A-D). To examine the distribution of mcherry-PopZ during the cell cycle, we performed a series of time-lapse microscopy experiments starting with a synchronized population of swarmer cells (Figure 4C). Images were acquired every one minute and kymographs were constructed to show the fluorescence intensity along the cell body over time. In wild-type cells, robust mCherry-PopZ foci accumulates at the new cell pole approximately 40 minutes post-synchrony (Figure 4C, movie 2). However, in a Δ*podJ* strain the vast majority of cells (90%, Figure 4D) proceed through cell division without accumulating any detectable PopZ at the new cell pole (Figure 4C, movie 3). Quantitative analyses in Figure 4D show that the proportion of the cells containing bipolar PopZ reaches the highest of ∼80% at 105-minute post-synchrony for wild-type and *podJ+* cells, comparing to the highest of 10% in Δ*podJ* cells. Collectively, these results show that PodJ is required for robust PopZ accumulation at the new cell pole in *C. crescentus*. A subpopulation that does accumulate at the new cell pole implicates redundant factors such as TipN(Laloux and Jacobs-Wagner, 2013) and ZitP(Berge et al., 2016) assist in PopZ recruitment at the new cell pole.

**Figure 4:**
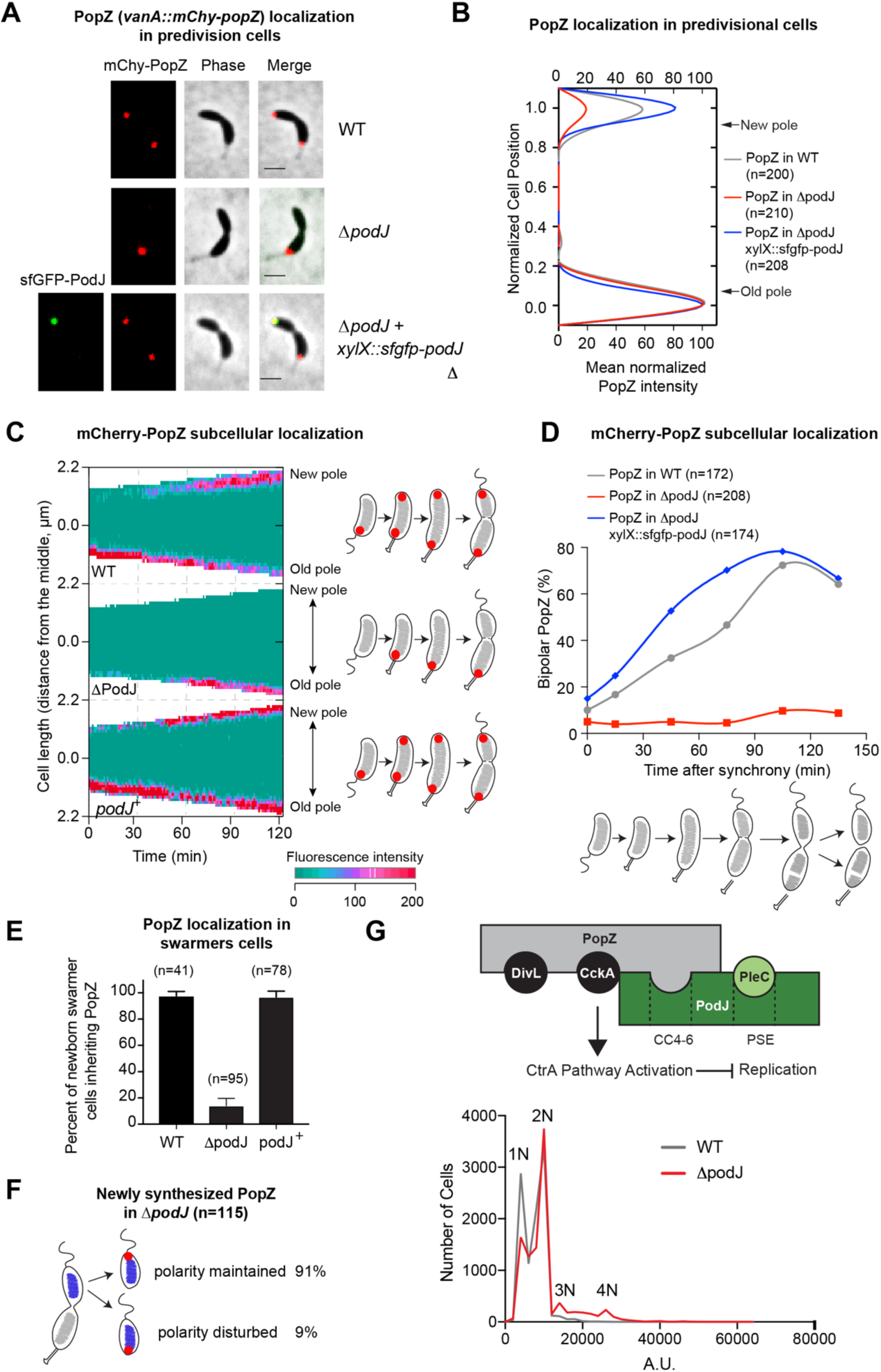
PodJ nucleates PopZ assembly at the new cell pole and promotes robust inheritance of PopZ in daughter swarmer cells. (A) mCherry-PopZ localization in predivisional cells in the wild-type (bipolar) versus the *podJ* deletion *C. crescentus* (monopolar). Bars, 2 μm. (B) Population analysis reveals a substantial reduction of PopZ abundance at the new cell pole of Δ*podJ* predivisional cells. Cell poles (new or old) were distinguished and orientated manually by observation of a stalk. The signal intensity was normalized with the highest value as 100% in each strain. (C) Kymograph analyses of mCherry-PopZ signal over the time after the synchronization of WT, Δ*podJ* and *podJ* complementary cells. Vanillate (50 mM) for mChy-PopZ induction and xylose (0.03%) for sfGFP-PodJ induction were added at 1 hr prior to synchronization of the cell culture. Images were acquired every 1 min. One of three representative cells were shown for each strain. The results indicate that PopZ fails to robustly accumulate at the new cell poles prior to cell division in Δ*podJ* strain. (D) Quantification of the percentage of cells that display detectable bipolar PopZ after cell synchronization. Time course analyses were performed within 135 mins in PYE medium for WT, rJ*podJ* and *podJ* complementary cells (more than 130 cells were calculated for each point). Robust PopZ assembly at the new cell pole is dependent upon PodJ. (E and F) Failure to localize mcherry-PopZ to the new cell pole in Δ*podJ* results in ∼80% population of swarmer cells that fail to inherit PopZ. Within a subpopulation of swarmer cells (9% of cells), nascent PopZ accumulates at the incorrect cell pole switching the inherited polarity axis. A total of 40 cells were tracked by time-lapse analyses. (G) Flow cytometry analysis of wild-type versus Δ*podJ* strains. When podJ is deleted from cells (red line), cells display a decrease in 1N and 2N cells that is accompanied with an increase in 3N and 4N cells.

### In the absence of PodJ, the cell polarity axis is randomized

In wild-type cells, PopZ accumulation at the new cell pole prior to cell division ensures that swarmer daughter cell inherit a PopZ signaling complex that maintains CtrA activation and prevents chromosome replication. In contrast, within the Δ*podJ* strain, the disability of PopZ to accumulate at the new cell pole led to 86% (82 out of 95 cells) of swarmer daughter cells void of PopZ (Figure 4E, Figure S4). In these cells, nascent PopZ is accumulated as monopolar focus approximately 60 minutes after cell division (Figure 4C). In *C. crescentus* Δ*podJ* strain, time-lapse analysis showed that in 91% of cells, PopZ accumulates at the same pole as in the wild-type strain (Figure 4F). However, in the other 9% of cells, PopZ accumulates at the opposite cell pole, disrupting the inherited cell polarity axis. These results indicate that PodJ is required to strictly maintain the inherited polarity axis for the newborn swarmer cells, and that other possible polarity factors (*e.g.*, SpmX or ParB) likely facilitate proper subcellular recruitment of nascent PopZ.

### CtrA pathway activation requires new cell pole assembly

Establishment of the new cell pole signaling complex is required for activation of the CtrA signaling pathway and repression of chromosome replication. Previous studies revealed that several new cell pole signaling proteins that promote CtrA phosphorylation (DivL, CckA, and PleC) displayed reduced accumulation at the new cell pole in the ΔPodJ strain(Curtis et al., 2012). Here we have shown that PodJ is a direct binding partner for PleC (Figure 3) and PopZ (Figure 4), which recruits the DivL/CckA complex(Holmes et al., 2016). To investigate the impact of new cell pole signaling hub disruption from the ΔPodJ strain, we examined the impact upon the CtrA signaling pathway activation by flow cytometry analysis of exponentially growing *WT* and Δ*podJ* cells stained with the nucleic acid dye SYTOX Green. Phosphorylated CtrA serves as a direct inhibitor of chromosome replication resulting in a tightly regulated cell population that contains either one or two chromosomes. A lower 1N:2N chromosome ratio was observed in Δ*podJ* cultures compared with *WT*, as well as a substantial increase in cells with three or more chromosomes (Figure 4G). These results indicate that PodJ’s scaffolding functions at the new cell pole promote CtrA signaling pathway activation.

### PopZ binds directly to coiled-coil region 4-6 of PodJ

To determine if PopZ and PodJ interact directly, we heterologously expressed fluorescent protein fusions of PodJ and PopZ in *E. coli*. As shown in Figure 5A and 5B, mCherry-PopZ accumulates at a single cell pole when it is expressed alone, while PopZ co-localizes in a bipolar pattern when co-expressed with YFP-PodJ. To determine the PopZ binding site within PodJ, we screened the capability of PopZ to bind to the library of PodJ domain deletion variants through co-expression in *E. coli* (Figure 5C, Figure S2). We used the following screening criteria to characterize PopZ interaction with the PodJ variants: (a) the two proteins are 100% co-localized and (b) the monopolar localization pattern of PopZ is changed after co-expression. We found that deletion of the C-terminal periplasmic domain or the intrinsically disordered PSE domain in PodJ did not disrupt the PodJ-PopZ interaction (Figure 5C, S2). In contrast, deletion of its CC4-6 domains disrupted PopZ co-localization with PodJ. We then expressed YFP-CC4-6 alone and observed that it was dispersed through the cytoplasm in *E. coli*. However, mCherry-PopZ was able to recruit it to the cell pole when co-expressed them in *E. coli.* These data indicate that coiled-coil 4-6 in PodJ functions as a PopZ recruitment site (Figure 5C).

**Figure 5:**
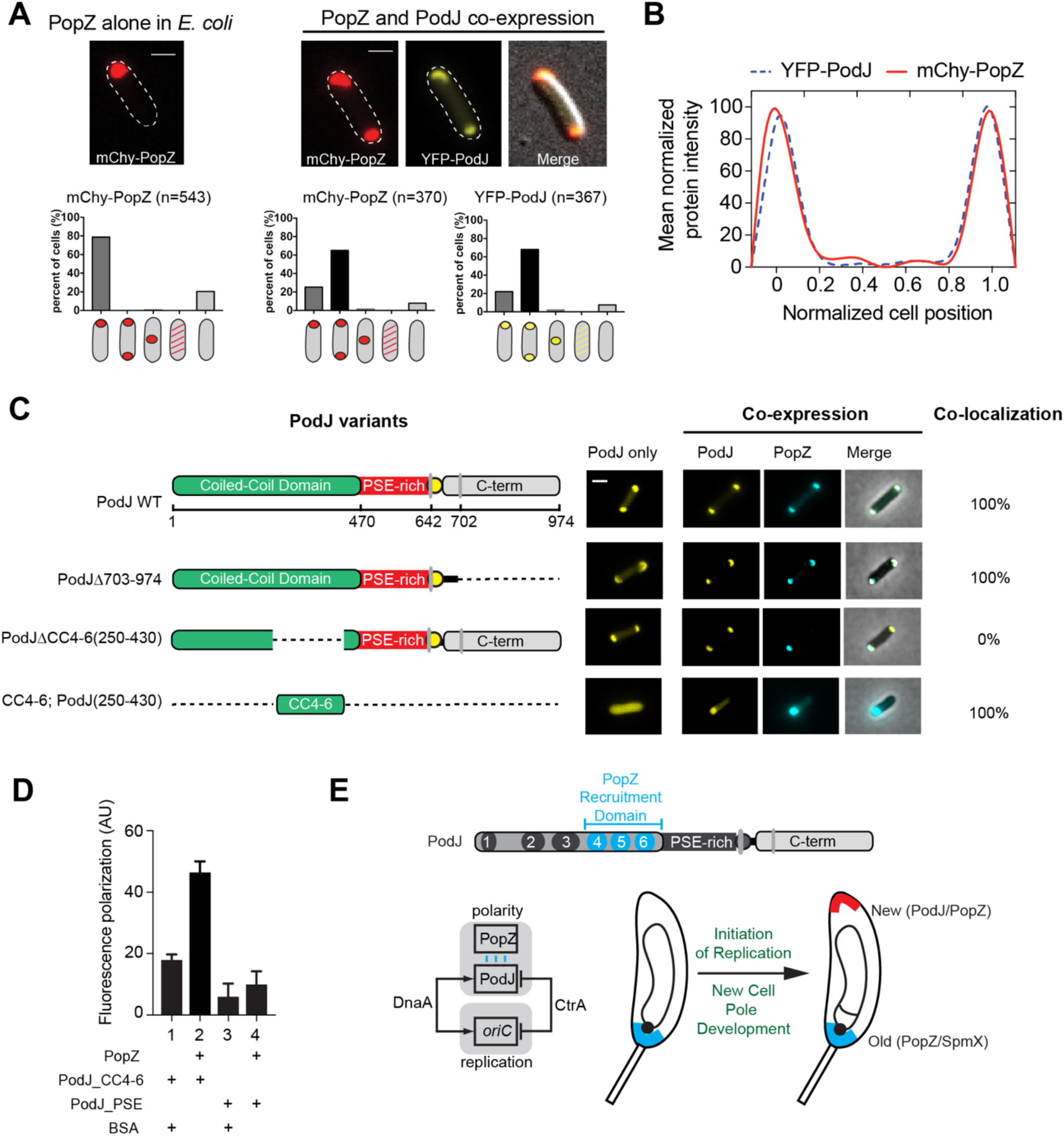
PopZ binds directly to the coiled-coil 4-6 region of PodJ. **(A)** Heterologous expression of YFP-PodJ and mcherry-PopZ in *E. coli*. Co-expression with PodJ causes bipolar PopZ accumulation in *E. coli*. (B) Mean protein intensity of YFP-PodJ and mcherry-PopZ versus cell length (n=370). The signal intensity was normalized with the highest value as 100% in each strain. (C) Co-expression of PodJ variants together with PopZ in *E. coli* reveals that the coiled-coil 4-6 region in PodJ is necessary for the interaction with PopZ. (D) Fluorescence polarization binding assay of the BODIPY dye labeled PodJ_PSE or PodJ_CC4-6 mixed with 10 µM PopZ, using BSA as a negative control. PopZ binds specifically to the CC4-6 domain of PodJ, however does not bind to its PSE-rich domain (E) Model for PodJ serving as a new cell pole development signal that triggers polarity establishment upon the initiation of replication, through its cell-cycle coordinated expression and specific interaction with the PopZ scaffold. All bars, 2 μm.

To confirm this PopZ-PodJ protein-protein interaction is direct, we purified the PodJ_CC4-6 protein with a cysteine incorporated after the hexahistidine purification tag and fluorescently labeled it as with PodJ_PSE. A fluorescence polarization assay was employed to detect a binding interaction between PodJ and PopZ by mixing 16 µM PopZ together with 100 nM BODIPY-PodJ_CC4-6, using the same amount of BODIPY-PodJ_PSE as a control. As shown in Figure 5D, PopZ bound to PodJ_CC4-6, but did not bind to PodJ_PSE. Taken together, the *E. coli* heterologous expression assays and *in vitro* biochemical assays show that coiled-coil 4-6 region of PodJ is the site of interaction with PopZ (Figure 5E). Moreover, we propose this PodJ-PopZ interaction plays a critical role in maturation of the new cell pole signaling hub (Figure 4A-F) and transcriptional regulation of PodJ(Crymes et al., 1999) coordinates this event with chromosomal replication (Figure 5E).

### SpmX is a negative regulator of PodJ accumulation at the cell poles

Our PodJ overexpression experiments (Figure 1D) and *E. coli* reconstitution experiments (Figure 2C) suggest that PodJ has an affinity for both cell poles. But how does PodJ exclusively bind and recognize the new cell pole in *C. crescentus*? Eukaryotic cell polarity networks are characterized by inhibitory regulatory interactions between the asymmetrically partitioned complexes to ensure robust cell polarization (Chau et al., 2012). Based upon these eukaryotic polarity networks, we hypothesized that proteins that reside at the old cell pole (*e.g.*, SpmX or DivJ) might play a role as a negative regulator of PodJ subcellular accumulation (Chau et al., 2012). We found that overexpression of SpmX resulted in a significant reduction of PodJ at the cell poles compared to that in wild-type cells (Figure 6A, 6C). In contrast, sfGFP-PodJ accumulated at all cell poles including constricted mid-cell curved sites in Δ*spmX* mutant strains (Figure 6B, 6C). These results suggest that SpmX is a negative regulator of PodJ accumulation at the cell poles either directly or indirectly.

**Figure 6:**
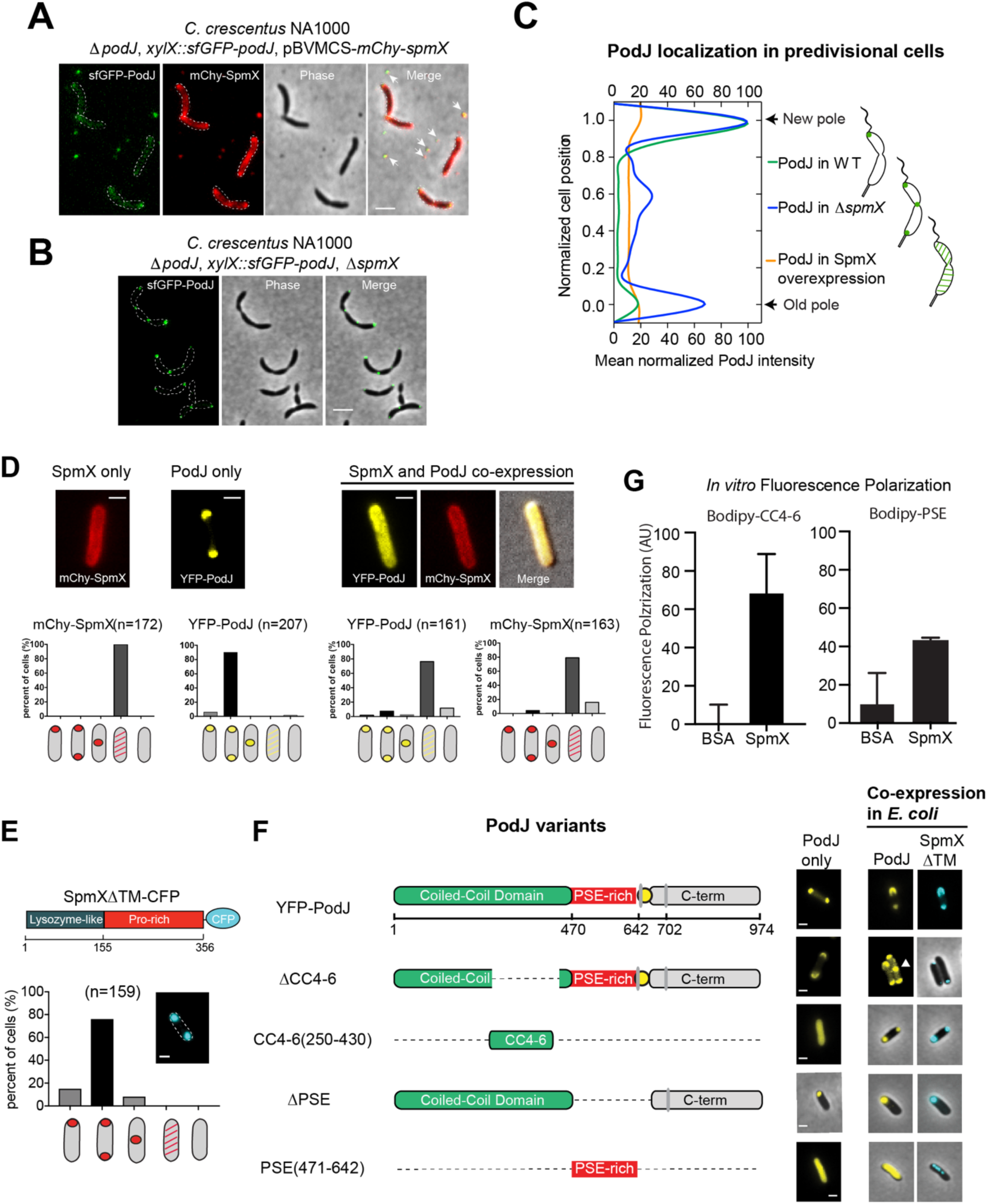
SpmX is a negative regulator for cell pole accumulation of PodJ. (A) SpmX directly or indirectly regulates PodJ subcellular localization in *C. crescentus*. Overexpression of SpmX results in a reduction of cell pole localized PodJ and secretion of PodJ from cells. (B) In the absence of SpmX, PodJ accumulates at all poles in *C. crescentus*. (C) Quantitative analysis of PodJ localization in *C. crescentus* predivisional cells in wild-type, Δ*spmX* and SpmX overexpression strains. (D) Heterologous expression of YFP-PodJ alone results in bipolar PodJ accumulation, while expression of SpmX-mcherry alone is disperse in *E. coli*. (E) Co-expression of PodJ together with SpmX causes PodJ to disperse from the cell poles in *E. coli*. All bars, 2 μm. (F) Co-expression of PodJ variants together with a SpmX variant lacking the transmembrane domain (SpmX’). These analyses indicate that SpmX’s dispersal of PodJ requires the PSE and CC4-6 domains of PodJ. (G) I*n vitro* fluorescence polarization assays screening the binding interactions between SpmX and Bodipy labeled PodJ-PSE domain and PodJ-CC4-6 domains.

We further examined if SpmX could disrupt PodJ subcellular accumulation at the cell poles when co-expressed in *E. coli* (Figure 6D). Consistent with previous studies (Holmes et al., 2016; Perez et al., 2017), SpmX expression alone is diffuse in *E. coli*. PodJ itself accumulates at the cell poles however co-expression of SpmX and PodJ together resulted in dispersion of the PodJ (Figure 6D). These results suggested that PodJ and SpmX interact directly and that dispersal of PodJ from the cell poles depends on the increase of SpmX protein level, consistent with the up-regulation of SpmX at the G1 transition phase in *C. crescentus* (Radhakrishnan et al., 2008; Schrader et al., 2016) (Figure 1A).

### SpmX interacts with the CC4-6 and PSE-rich domains of PodJ

To determine the SpmX interaction domain, we co-expressed a library of PodJ domain deletions in *E. coli* with a SpmX variant lacking the transmembrane domains, hereafter called SpmX’. While full-length SpmX is disperse throughout the cytoplasm, SpmX’ accumulates at both cell poles. We observed that co-expression of PodJ with SpmX’ resulted in PodJΔCC4-6 foci accumulation away from the cell poles. When the PodJ CC4-6 variant is expressed by itself in *E. coli* it was diffuse (Figure 6E, 6F), however SpmX’; is capable of recruiting PodJ CC4-6 to the cell poles. This result indicates that SpmX’; binds directly to coiled-coil 4-6 region of PodJ. We then more closely examined the role of PodJ’;s PSE domain in the PodJ-SpmX interaction. Upon SpmX co-expression with PodJΔPSE we observed that the two proteins co-localized at the cell poles without any observed dispersal from the cell poles. We also observed that expression of the PSE-rich domain by itself could disperse SpmX from the cell poles in *E. coli*. These data indicate that SpmX dependent dispersion of PodJ also requires the PSE domains. As the CC4-6 and PSE domains are adjacent to one another they may work collectively to regulate the PodJ-SpmX interaction.

To confirm PodJ and SpmX interact directly, we purified SpmX’, and employed the fluorescence polarization assay to detect the interaction between SpmX’ and the BODIPY-PodJ_CC4-6 or BODIPY-PodJ_PSE. As shown in Figure 6G, SpmX’ interacted with both BODIPY-PodJ_CC4-6 and BODIPY-PodJ_PSE. Interestingly, the consequences of these two binding interactions are distinct: SpmX binding to CC4-6 region results in co-localization, while SpmX binding to the PSE-domain results in PodJ and SpmX dispersal from the cell poles.

### PodJ interactions with PopZ and SpmX are required for robust cell polarity

Our experiments suggest a 3-node protein-protein interaction circuit in which PodJ promotes PopZ new cell pole accumulation (Figure 4), PopZ promotes SpmX subcellular accumulation(Perez et al., 2017), and SpmX negatively regulates PodJ subcellular accumulation (Figure 6). We introduced PodJ variants as a sole copy into *C. crescentus* that would individually disrupt the proposed regulatory interactions and evaluated their impact upon *C. crescentus* cell polarity: PodJΔperi, PodJΔCC4-6 and PodJΔPSE. When sfGFP-PodJΔPSE is expressed as a sole copy in *C. crescentus*, we observed that sfGFP-PodJΔPSE and mCherry-PopZ accumulates at both cell poles in predivisional cells (Figure 7A). This indicates the PSE domain in PodJ, the site of PodJ-SpmX interaction, is critical for PodJ to be excluded from the old cell pole. When sfGFP-PodJΔCC4-6 is expressed as a sole copy in *C. crescentus*, we found that PopZ-mCherry is poorly localized at the new cell pole (Figure 7A), similar to the observation that was found in Δ*podJ* (Figure 4B). Therefore, deletion of CC4-6 disrupts polarity through loss of PopZ recruitment at the new cell pole. In contrast, sfGFP-PodJΔperi construct mirrors wild-type like cell polarity (Figure 7A) as it contains critical regulatory interactions with PopZ and SpmX.

**Figure 7:**
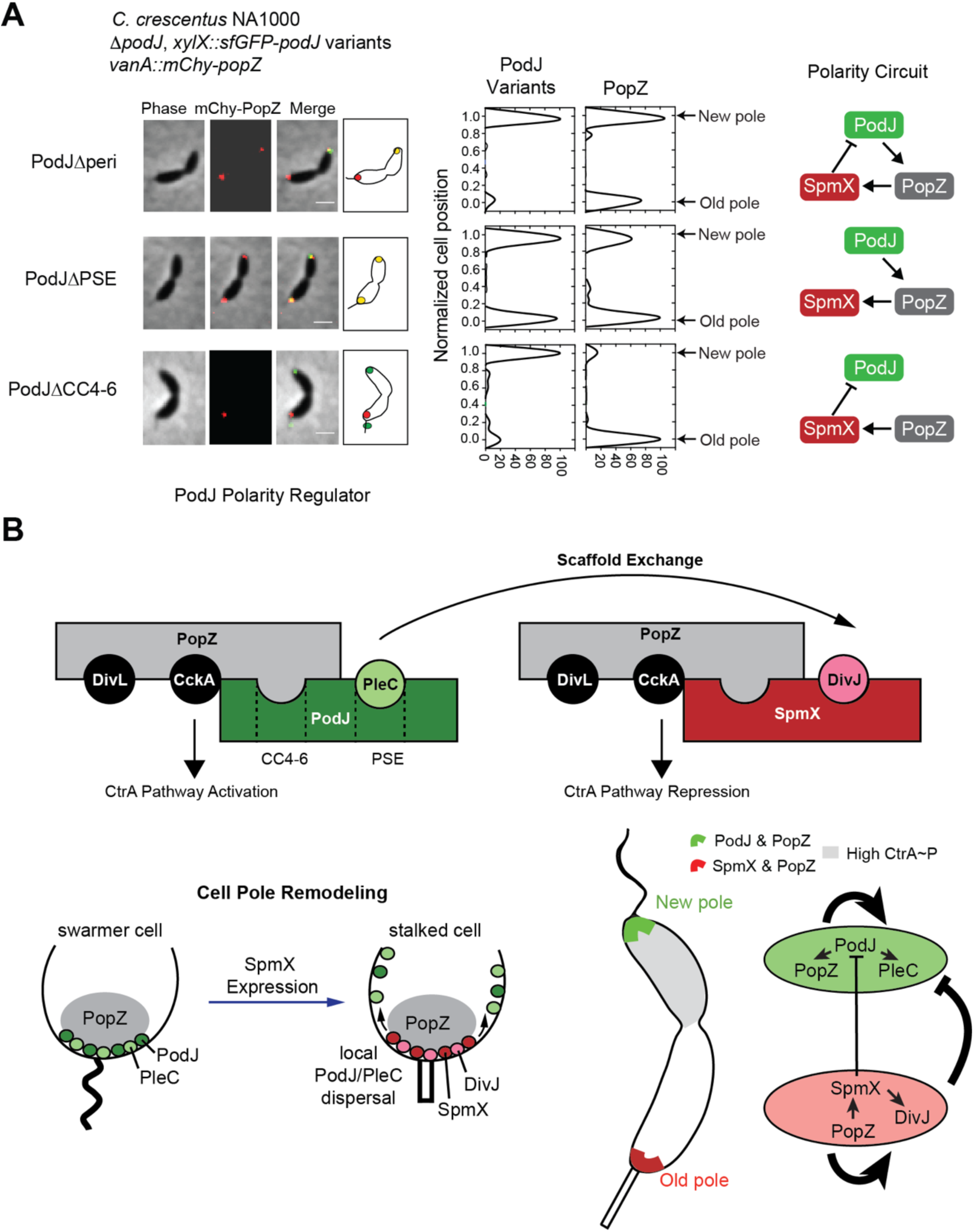
A network of protein-protein regulatory interactions enable polarity establishment in *C. crescentus*. (A) A polarity circuit in which PodJ promotes PopZ new cell pole localization, PopZ promotes SpmX accumulation at the cell poles, and SpmX negatively regulates PodJ subcellular localization at the cell poles Expression of PodJ variants and PopZ as the sole copy in *C. crescentus*. The cytoplasmic domain of PodJ, containing both CC4-6 and the PSE-rich domain, supports bipolar localization of PopZ and new cell pole localization of PopZ. In contrast, expression of PodJΔCC4-6 results in predivisional cells that fails to accumulate PopZ at the new cell pole. Removal of the domain that is required for SpmX to disperse PodJ from the cell poles, PodJΔPSE, results in PodJ accumulation at both cell poles in *C. crescentus*. These results indicate that the regulatory protein-protein interaction between PodJ, PopZ and SpmX are required for robust cell polarity of *C. crescentus* predivisional cells. **(B)** SpmX functions as a local negative regulator of PodJ accumulation at the old cell pole. **(C)** The establishment of the compositionally distinct new and old cell poles is driven by a network of scaffolding proteins that combines positive feedback together with inhibition.

To our surprise, we observed that sfGFP-PodJΔCC4-6 is secreted from the stalked pole in wild-type cells (Figure S7, Figure 7A). However, this abnormal phenotype can be restored by deleting *spmX* (Figure S7). Moreover, besides dispersal of intracellular PodJ, overexpression of SpmX also resulted in secretion of PodJ foci from the cell (Figure 6A, Movie S4), indicating SpmX plays a role in secretion of PodJ. Similar secretion phenotype was observed when expression sfGFP-PodJ after deletion of PopZ (Figure S7), suggesting that PopZ anchors PodJ in the cytoplasm and prevents this abnormal secretion process. To exclude the possibility that PodJ is excreted due to cell lysis, we confirmed sfGFP-PodJ by western blot analyses of the media supernatants with anti-GFP antibody, using the intracellular CtrA protein as a negative control (Figure S7). These results collectively suggest that SpmX plays a negative regulatory role in dispersing PodJ from the old cell pole, a function that we propose is critical for the new-to-old cell pole remodeling that triggers swarmer-to-stalk cell differentiation (Figure 7B).

## Discussion

Here we have shown that the protein-protein interactions between three scaffolding proteins function together as a circuit to establish and maintain polarity in *C. crescentus.* Using a synthetic biology reconstitution approach, we have identified the localization hierarchy at the new cell pole by demonstrating that PodJ recruits three client proteins (PleC, CpaE, PopA), and the central organizing scaffold PopZ. In turn, this allows PopZ to recruit its direct binding partners (DivL, CckA) to the new cell pole (Figure 3F) (Holmes et al., 2016). Thus the PodJ scaffold promotes the colocalization of PleC together with the PopZ/DivL/CckA signaling complex to promote CtrA pathway activation (Figure 4G). To trigger asymmetric accumulation of PodJ, our studies have also revealed that the old cell pole scaffold SpmX promotes localized foci disassembly and an unexpected stalked pole specific secretion process (Figure 6-7). Taken together we propose that this scaffold network topology composed of both positive feedback and negative regulations between signaling hubs promotes robust cell polarization (Figure 7).

### PodJ serves as a central node in the new cell pole signaling hub localization hierarchy

Prior to this study it was known that several factors accumulate at the new cell pole (Curtis and Brun, 2010; Lasker et al., 2016), however the localization dependency within this new cell pole signaling hub was unclear. We confirmed that three protein-protein interactions with the client proteins CpaE, PleC and PopA were direct. Most critically, we discovered a direct PodJ-PopZ interaction that further triggers accumulation of PopZ binding proteins (DivL and CckA). In the absence of PodJ, the PopZ scaffold still can accumulates at the new cell pole in a fraction of cells (Figure 4), consistent with contribution of other redundant factors such as ZitP(Berge et al., 2016) and TipN(Laloux and Jacobs-Wagner, 2013). These studies indicate that the PodJ scaffold serves as a central organizer of the new cell pole development (Figure 3F). Key questions remain as to the mechanism of how this nucleating factor targets the cell poles? Our studies indicate that oligomerization of PodJ (Figure 2) may allow PodJ to recognize micron sized curvature differences (Huang and Ramamurthi, 2010) or interact with shared cell wall components between *C. crescentus* and *E. coli* such as the Tol-Pal membrane integrity complex (Yeh et al., 2010), pili and flagellar assemblies.

### SpmX prevents symmetric PodJ localization

A key aspect of eukaryotic cell polarity circuits are regulatory interactions between partitioned complexes(Chau et al., 2012). Our studies have revealed that SpmX is a novel negative regulator of PodJ localization. Deletion of SpmX results in accumulation of PodJ at all highly curved regions of the cell (Figure 6A) while overexpression of SpmX results in dispersion of PodJ from the cell poles and an unexpected localized secretion of PodJ at the stalked cell pole (Figure 6A, Figure S7(Grangeon et al., 2015)). We propose that this local PodJ dispersal functions may play a role in compositional remodeling of the PodJ-rich signaling hub into a SpmX-rich signaling hub that occurs in the swarmer-to-stalked cell differentiation (Figure 7B).

PodJ domain analysis indicates that SpmX binds to two PodJ domains: CC4-6 and the instrically disordered PSE-rich domain. The biophysical mechanism of this foci disassembly process remains unclear, and one possible model is that PodJ-SpmX complex forms small oligomers incapable of cell pole recognition. We speculate that the SpmX-PodJ regulatory interactions may share broad mechanistic similarities with the MipZ-FtsZ (Thanbichler and Shapiro, 2006) and (MinD-FtsZ) (Park et al., 2018) interactions that prevent FtsZ accumulation at the cell poles and direct FtsZ to the mid-plane. However, extensive future *in vitro* studies will be needed to fully understand this dispersal mechanism. As well, we anticipate that SpmX likely mediates other cell pole remodeling mechanisms indirectly which will require future genetic studies to identify new regulatory partners and interactions. It also remains to be seen if co-localization of PodJ, the ClpXP adaptor protein PopA and ClpXP promote the final stages of PodJ_S_ proteolysis at the old cell pole to promote asymmetric localization.

### Cell polarity is achieved though polarity circuits with common network topology throughout all kingdoms of life

The PopZ-SpmX-PodJ polarity circuit is conserved within a subset of α-proteobacteria, and recent studies of have revealed genetic interactions between PodJ and PopZ in *Agrobacterium tumefaciens* (Anderson-Furgeson et al., 2016; Grangeon et al., 2015), however the subcellular localization pattern of these proteins diverges from that observed in *C. crescentus* (Anderson-Furgeson et al., 2016; Ehrle et al., 2017; Grangeon et al., 2015; Howell et al., 2017). In *A. tumefaciens*, the polarity is inverted compare to *C. crescentus* as PopZ residues exclusively at the new cell pole and PodJ occupies the old cell pole (Grangeon et al., 2015). Moreover, a second subset of α-proteobacteria species encode the PopZ scaffolding protein and the PleC and CckA histidine kinases, however their genomes contain no clear homologs of the PodJ or SpmX scaffolding proteins. These variations in conservation suggest that that the PopZ-SpmX-PodJ polarity network may have been re-wired to include new regulatory interactions and new regulators to support diverse modes of bacterial cell development observed throughout α-proteobacteria (Ettema and Andersson, 2009).

More broadly, polarity networks are remarkably diverse at the biomolecular level and are utilized throughout all kingdoms of life. *C. crescentus* orchestrates cell polarity through a network architecture that contains positive feedback through formation self-assembled scaffolding proteins. With positive feedback alone, PodJ self-assembles or binds to cell pole binding sites nearly equally well at each cell pole. In this study we have critically examined the inter-relationships between three scaffolds that promote *C. crescentus* cell polarity. Based upon these studies we propose a model that compositional control of the new and old cell pole signaling hubs is promoted by local disassembly of PodJ at the old cell pole by a PodJ foci inhibitor protein SpmX (Figure 7B). Remarkably, this network topology of positive feedback coupled with inhibition is similar to minimal polarity circuit architectures identified using a synthetic biology approach in yeast (Chau et al., 2012). This synthetic biology approach characterizing several network architectures suggested that mutual inhibition networks yielded the most robust cell polarization (Chau et al., 2012). Therefore, we anticipate future research will identify several negative regulatory interactions between the new and old cell pole signaling hubs to ensure robust *C. crescentus* cell polarization.

## Acknowledgements

We thank Jared Schrader for providing critical reviews of the manuscript. We also thank Lucy Shapiro for providing critical *C. crescentus* strains that supported this study.

## METHODS

### Phase Contrast, DIC, and Epifluorescence Microscopy

Cells were imaged after being immobilized on a 1.5% agarose pad containing corresponding inducers when required. Phase microscopy was performed by using a Nikon Eclipse T*i*-E Inverted microscope equipped with an Andor Ixon Ultra DU897 EMCCD camera and a Nikon CFI Plan-Apochromat 100X/1.45 Oil objective. DIC microscopy was performed using the same microscope and camera but with a Nikon CFI Plan-Apochromat 100X/1.45 Oil DIC objective with a Nikon DIC polarizer and slider in place. Excitation source was a Lumencor SpectraX light engine. Chroma filter cube CFP/YFP/MCHRY MTD TI was used to image ECFP (465/25M), EYFP (545/30M), and mCherry (630/60M). Chroma filter was used to image EGFP and sfGFP (470/40X, 515/30M). Chromosomal DNA was visualized by using 1.5 µg/ml DAPI. DAPI was imaged using 395/25X, 435/26M Chroma filter set. Images were collected and processed with Nikon NIS-Elements AR software.

### Time-lapse Microscopy

sfGFP-PodJ, mcherry-PopZ, SpmX-mcherry, or CFP-ParB was tracked using phase and fluorescence microscopy. Images were collected every 1-3 min over the course of 1-2 cell division (∼4 h). The imaging system used was Nikon Eclipse T*i*-E microscope equipped with an Andor Ixon Ultra DU897 EMCCD camera and NIS-Elements software. *C. crescentus* cells with corresponding expression gene were grown to the early-log phase in M2G or PYE medium (OD_600_ = 0.2), and then induced by xylose or vanillic acid for 2 hours prior to synchronization. Swarmer cells were isolated from the culture by centrifugation (20 mins at 11,000 rpm, 4°C) after mixture with 1 volume of Percoll (GE Healthcare). The synchronized swarmer cells were pipetted onto an agarose (2%) pad containing medium with inducers, and sealed with wax. During time lapse experiments, phase and fluorescence images were taken in 1 min intervals for sfGFP-PodJ, mcherry-PopZ, and SpmX-mcherry. Phase and fluorescence images were taken in 3 min intervals for CFP-ParB.

### Purification of PodJ, PopZ, SpmX, and PleC Variants

Protein expression of all PodJ variants followed the same protocol and is described in detail below for the PodJ (1-635). To purify the cytoplasmic portion of PodJ(1-635), Rosetta (DE3) containing plasmid pwz091 was grown in 6 liters LB medium (20 µg/ml chloramphenicol and 100 µg/ml ampicillin) at 37°C. The culture was then induced at an OD600 of 0.4–0.6 with 0.5 mM IPTG overnight at 18°C. The cells were harvested, resuspended in the lysis buffer (50 mM Tris-HCl, 700 mM KCl, 20 mM Imidazole, 0.05% dextran sulfate, pH 8.0), in the presence of protease inhibitor cocktail tablets without EDTA (Roche).

The cell suspension was lysed with three passes through an EmulsiFlex-C5 cell disruptor (AVESTIN, Inc., Ottawa, Canada), and the supernatant was collected by centrifuging at 13000 *g* for 30 min at 4°C. In addition, the insoluble cell debris was resuspended by the recover buffer (50 mM Tris-HCl, 1000 mM KCl, 20 mM Imidazole, 0.05% dextran sulfate, pH 8.0) and its supernatant was collected as well as the previous centrifugation. The combined supernatants were loaded onto a 5 ml HisTrapTM HP column (GE Healthcare) and purified with the ÄKTA™ FPLC System. After washing with 10 volumes of wash buffer (50 mM Tris-HCl, 300 mM KCl, and 25 mM imidazole, pH 8.0), protein was collected by elution from the system with elution buffer (50 mM Tris-HCl, 300 mM KCl, and 500 mM imidazole, pH 8.0), and concentrated to a 3 ml volume using Amicon Centrifugal Filter Unites, resulting in > 95% purity. All PodJ variants were dialyzed with a buffer containing 50 mM Tris-HCl (pH 8.0), 300 mM KCl, and then aliquoted to small volume (100 µl) and kept frozen at −80°C till use.

His-SpmX (1-365) REF and His-PleC_PASCD expression and purification followed the same protocol except using a different lysis buffer (50 mM Tris-HCl, 300 mM KCl, 20 mM Imidazole, pH 8.0) and without recover step as in PodJ purification. His-PopZ was expressed and purified the same as described (Ptacin et al., 2014).

### Size Exclusion Chromatography and Native Gel Assay

A gel filtration standard (Sigma) containing thyroglobulin (bovine, 669 kDa), carbonic anhydrase (bovine, 29 kDa), blue dextran (2,000 kDa), apoferritin (horse, 443 kDa), β-Amylase (sweet potato, 200 kDa), alcohol dehydrogenase (yeast, 150 kDa), and albumin (bovine, 66 kDa) were used to generate a molecular weight standard plot using a Superdex 200 10/300 GL column (GE Healthcare). A 3.2 mg/ml sample of His-PodJ(1-635) was loaded onto the column and eluted after 7.9 ml, 12.8 ml, and 15.0 ml of buffer, corresponding to a molecular weight of 1,851 kDa, 194 kDa, and 70.7 kDa (theoretical monomer = 73.0 kDa). One representative result of triplicates was shown.

His-PodJ(1-635) was also analyzed by running a native gel. Protein was separated by gel electrophoresis (8% separate gel) at 80 V for at least 4 hours at 4oC (ref), using a Native protein ladder (range from 66 to 669 kDa, Thermo Fisher).

### Fluorescence Polarization Assay

To label PodJ_PSE (471-635) and PodJ_CC4-6 (250-430), we cloned a cysteine just after the 6X His tag of these two proteins at the N-terminal. PodJ_PSE (Cys) and PodJ_CC4-6 (Cys) expression and purification followed the same protocol as PodJ mentioned above. These two proteins were labeled at the cysteine using thiol-reactive BODIPY™ FL N-(2-Aminoethyl) Maleimide (Thermo Fisher). The proteins were mixed together with 10-fold excess BODIPY™ FL N-(2-Aminoethyl) Maleimide and allowed to react for 2 hours at room temperature, and the unreacted dye was quenched with mercaptoethanol (5% final concentration). The labelled proteins were purified via dialysis to remove unreacted fluorescent dye (5 times, 500 ml buffer and 30 mins each).

Fluorescence polarization binding assays were performed by mixing 100 nM labeled proteins with 0, 0.25, 0.5, 1, 2, 4, 8, 16 µM partner protein (PopZ, SpmX, PleC, or BSA) for 45 minutes to reach binding equilibrium at the room temperature. Fluorescent Proteins were excited at 470 nm and emission polarization was measured at 530 nm in an UV-vis Evol 600 spectrophotometer (Thermo). Fluorescent polarization measurements were performed in triplicates, and three independent trials were averaged with error bars representing the standard deviation.

### Western Blot

Western blot analysis was used to determine if PodJ was excreted from cells upon overexpression of SpmX and in Δ*popZ* strains. Cells were grown in 30 ml PYE medium to early-log phase and induced by appropriate inducer for 3 hours at 30°C. Cells were harvested at OD600 about 0.6 (4,000 *g*, 10 min) and resuspended using 5 ml PYE medium, and the supernatants were also collected. Next, cells were removed from supernatants by filtration using a 0.45 µm pore membrane (GE Healthcare). The supernatants containing excreted PodJ were then condensed into 1 ml by additional centrifugation (30,000 *g*, 60 min). The presence of purified PodJ foci from media supernatants were confirmed by observing PodJ foci using epifluorescence microscopy.

Both cells and their supernatants were lysed by heating (100°C for 30 min) with protein sample buffer. Approximately 5-µg protein from cell samples and the same volume samples from supernatants were loaded on and separated by 12% SDS–PAGE. Proteins were transferred onto PVDF membranes (GE Healthcare) and standard western blotting procedures (Li et al., 2015) were followed. The anti-GFP antibody (Cell Signaling Technology) and anti-CtrA antibody (gift from Lucy Shapiro) with 1:1000 dilution was used to determine the distribution of sfGFP-PodJ and CtrA in/out of the cell, respectively. PVDF membranes were treated with an ECL western blotting kit (ThermoFisher) and visualized using a ChemiDoc XRS+ system (Bio-Rad).

### Flow Cytometry

Strains analyzed were grown overnight in PYE under antibiotic selection pressure at 28°C. Cells were diluted to an OD_600_ of 0.1 in PYE/antibiotic and induced with 0.03% xylose for 4 hours, if applicable. Rifampicin (20 µg/mL) was then added and cells were grown for 3 more hours to allow for complete replication of DNA. Cells were then fixed in cold, 70% ethanol overnight at 4°C for up to one week by adding 700 µL of 200 proof ethanol to 300 µL of cell culture. To stain, cells were collected by centrifugation at 6,000g for 2 minutes and resuspended in 1mL of Tris-HCl buffer (20mM Tris-HCl, pH=7.5, 150mM NaCl) containing 0.2 µg/mL RNase (RNase A, 10 mg/mL, ThermoFisher) and 1µM SYTOX Green nucleic acid stain. Cells were incubated at room temperature for 15 minutes then samples were run on a Cytoflex S (Beckman-Coulter) using 488 nm laser with a FitC filter (525 nm). Cells were selected using FSC-H and SSC-H gained to 10 and 20, respectively. Cells were thresholded in FitC-H at 1000. 10000 events were collected flowing at 10 µL per minute with an abort rate of less than 5%. Raw data was exported to Prism and histograms were generated.

## QUANTIFICATION AND STATISTICAL ANALYSIS

FIJI/ImageJ ((Schindelin et al., 2012);(Preibisch et al., 2009)) and MicrobeJ (Ducret et al., 2016) were used for image analysis. Number of replicates and number of cells analyzed per replicate are specified in corresponding legends. All experiments were replicated at least 2 times, and statistical comparisons were carried out using GraphPad Prism with two-tailed Student’s t tests. Differences were considered to be significant when *p* values were below 0.05. In all figures, measurements are shown as mean ± standard deviations (s.d.).

### Kymograph Analyses

Kymographs of fluorescence intensity were obtained by using the built-in kymograph function of the Microbe-Tracker in MicrobeJ(Ducret et al., 2016). The signal of background was subtracted before the kymograph analysis, and the observation of stalk at the pole of *C. crescentus* cell was defined as the old pole. The predivisional cell was selected as the start point cells in Figure 1C and Figure 4G. In Figure 1C, another round of kymograph analysis was performed after the first cell division. The new pole **b** became to be the old pole after cell division and another two new pole (**c** and **d**) were formed.

### Intrinsically Disordered Region Analysis

The probability of intrinsic disorder region over the primary sequence of PodJ was predicted by three independent programs, *i.e*., Metadisorder MD2(Kozlowski and Bujnicki, 2012) SPOT (Hanson et al., 2017), and Cspritz(Walsh et al., 2011). The average scores of these programs were plotted against the PodJ sequence. We assumed the region as an intrinsic disorder with the probability > 75% in this study.

### Recruitment index measurement

The recruitment index in Figure 3B is calculated using the formula below similar as previous described(Holmes et al., 2016).

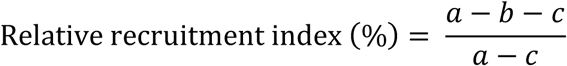

Here, *a* is the sum of the fluorescent signal within all the cell meshes, *b* is the sum of the fluorescent signal in the middle of cells, and *c* is the background fluorescence.

### Calculation of Subcellular Co-Localization with PodJ variants

To measurement of the co-localization ratio in Figure 3C, Figure 5C, Figure 6C, and Figure S3, we used strict criteria to calculate how the proteins interaction with the PodJ variants, *i.e.,* (I), the localization patterns of the interaction proteins are changed after co-expression. (II), the two proteins are 100% co-localized at the pole (binding), or drive each other apart from the pole (dispersion). Failure of either of these two criteria means the interaction of the two proteins is undetermined. About 200 cells were calculated for each interaction sets.

## Supplementary Figure Captions

**Figure S1:** (A) Time-lapse imaging of sfGFP-PodJ induced by 0.003% xylose after two rounds of cell division. (B) sfGFP-PodJ accumulates at the new cell pole in stalk and predivisional cells. During the swarmer to stalked cell transition sfGFP-PodJ diminishes at the old cell pole.

**Figure S2:** Analysis of PodJ domain deletion library when expressed alone heterologously in *E. coli* or co-expressed with PopZ, SpmX or PleC fluorescent protein fusions. Solid circles indicate co-localization of PodJ variants together with PopZ, SpmX or PleC. Open circles indicate PodJ variants that do not co-localize with PopZ, SpmX or PleC. Question marks indicate no assignment can be made based upon the co-expression assay.

**Figure S3:** (A) Analysis table of PodJ co-expression with potential client proteins in *E. coli*. (B) Co-expression of YFP-PodJ together with inclusion body protein A (IbpA-mChy) indicates that PodJ does not co-localize with inclusion bodies in *E. coli.* (C) Three potential PodJ protein-protein interaction partners (TipN, FtsZ and SpmX) promoted dispersion of YFP-PodJ when co-expressed in *E. coli.* (D) Co-expression of YFP-PodJ together with new cell pole associated proteins (PleD, DivL, DivK, CckA) indicate that these proteins do not co-localize when expressed in *E. coli.*

**Figure S4:** Subcellular localization pattern of *Ppopz-mCherry-popZ* in wild-type, *ΔpodJ*, and *ΔpodJ xylX::sfGFP-podJ* strains in the presence of 0.03% xylose in *C. crescentus* (A) pre-divisional cells and (B) newborn swarmer cells just after cell division. (C) Expression of mcherry-PopZ in BL21 *E. coli* indicates that PopZ accumulates randomly as a single focus at either the old or new cell poles.

**Figure S5:** Heterologous co-expression of PleC together with 3 new cell-pole associated scaffolds (PopZ, PodJ, and TipN). These assays indicate that PleC can be directly recruited to the cell pole by PodJ, while PleC is indirectly associated with the PopZ and TipN scaffold proteins.

**Figure S6:** (A) SpmX domain deletion library when expressed alone heterologously in *E. coli,* or co-expressed with PodJ fluorescent protein fusions. Dispersion of YFP-PodJ from the cell pole binding site requires the transmembrane domain of SpmX. The N-terminal fluorescent protein fusion of SpmX(1-356) disrupts its capability to accumulate as a focus suggesting that the N-terminus of SpmX may be involved in self-assembly. The SpmX-PodJ interaction requires both the lysozyme and proline-rich domains of SpmX. (B) Select PodJ domain deletion library variants when expressed alone heterologously in *E. coli* or co-expressed with SpmX or SpmXΔTM fluorescent protein fusions. These results suggest that PodJ’;s PSE and CC4-6 domains are sites of interaction with SpmX.

**Figure S7.** The PopZ-PodJ interaction anchors PodJ in the cytoplasm and prevents PodJ cellular secretion. (A) PodJ is specifically secreted from *C. crescentus* strains that disrupt the PodJ-PopZ interaction (PodJ ΔCC4-6), and (B) the PodJ secretion of C. crescentus requires the SpmX scaffolding protein. (C and D) Full-length sfGFP-PodJ is not secreted from wild-type cells, but is secreted from cells in the PopZ deletion strain (ΔPopZ). (E) Fractionated media indicates the presence of sfGFP-PodJ foci in growth media strains that trigger PodJ cellular secretion. (F) Western blot of analysis of sfGFP-PodJ and CtrA inside (I) and outside (O) of the cell. The ΔPopZ strain exhibits the largest amount of extracellular sfGFP-PodJ, while western blot controls of CtrA indicates that global cell lysis is likely not contributing to the observed PodJ secreted foci.

Movie S1: Time-lapse imaging of sfGFP-PodJ in *C. crescentus NA1000.*

Movie S2. Time-lapse imaging of mcherry-PopZ in *C. crescentus NA1000.*

Movie S3: Time-lapse imaging of mcherry-PopZ in the *C. crescentus ΔpodJ strain.*

Movie S4: Time-lapse imaging of sfGFP-PodJ cellular secretion upon SpmX overexpression.

