## Supplemental Information for "A circuit of protein-protein regulatory interactions enables polarity establishment in a bacterium"

#### Supplemental Figures

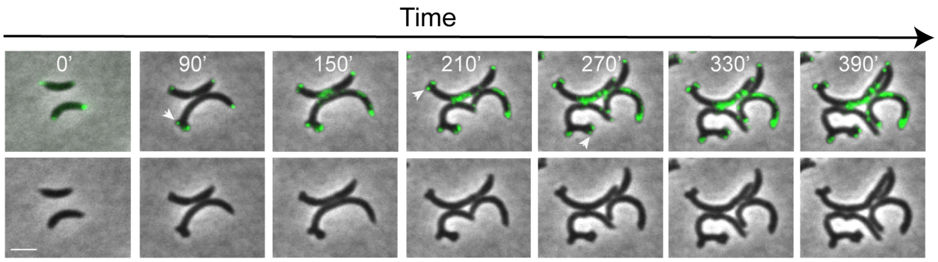


Figure S1: Time-lapse imaging of PodJ overexpression (0.3% xylose) in *C. crescentus.*

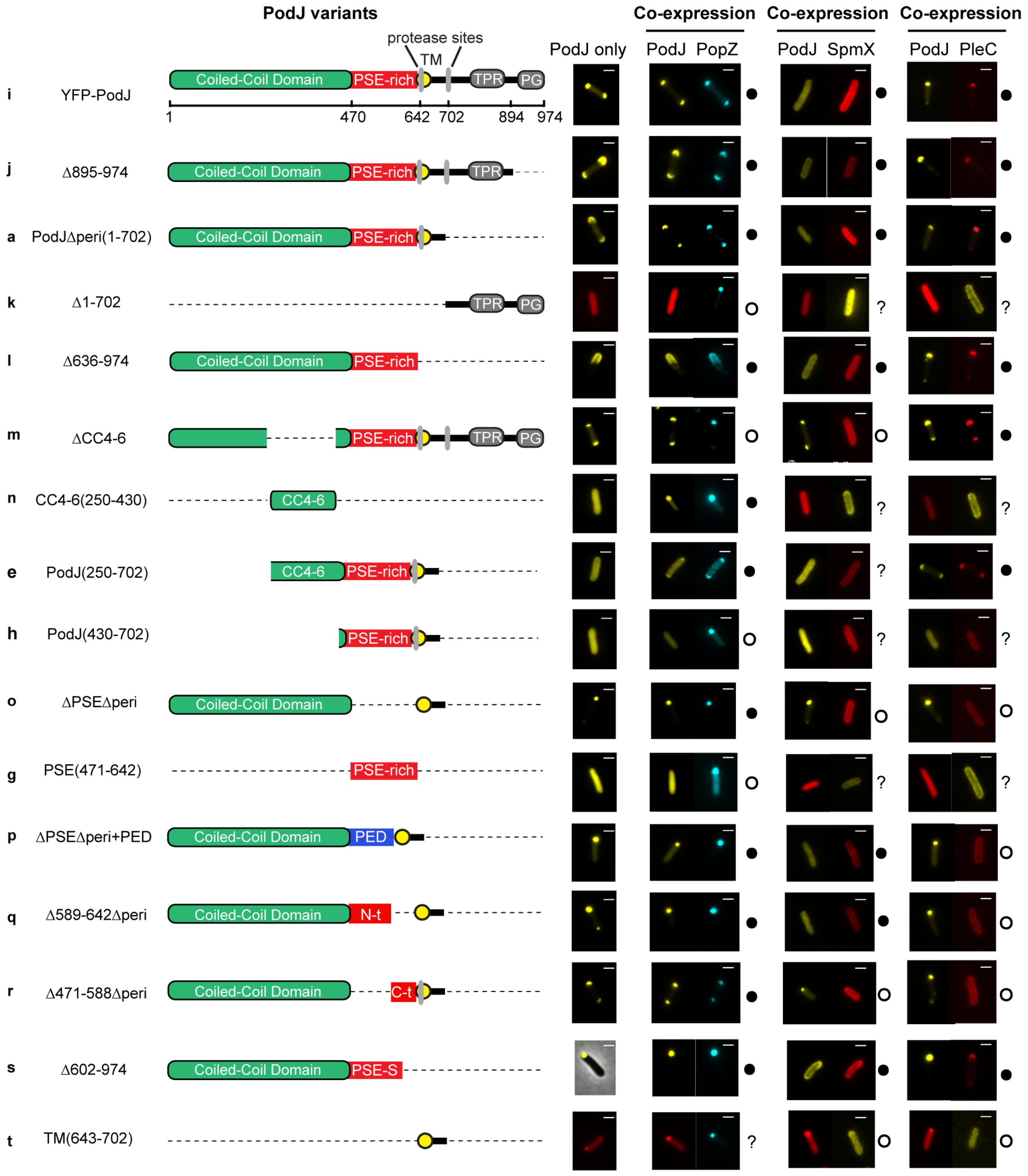


Figure S2: Analysis of PodJ domain deletion library when expressed alone heterologously in *E. coli,* or co-expressed with PopZ, SpmX or PleC fluorescent protein fusions. Solid circles indicate co-localization of PodJ variants together with PopZ, SpmX or PleC. Open circles indicate PodJ variants do not co-localize with PopZ, SpmX or PleC. Question marks indicate no assignment can be made based upon the current co-expression assay.

**A**

| Protein ID | Proteins | Description | Ortholog in *E.coli* BL21 (Identity) | Localization in *E. coli* | | |
|---|---|---|---|---|---|---|
| | | | | Alone | With YFP-PodJ or CFP-PodJ | With mChy-PopZ or CFP-PopZ |
| CCNA_02125 | YFP-PodJ | Scaffolding | ND | Bipolar | / | Co-localized, bipolar |
| **CCNA_01380** | **mChy-PopZ** | **Scaffolding** | **ND** | **Monopolar** | **Co-localized, bipolar** | **/** |
| **CCNA_01552** | **TipN-mChy** | **Scaffolding** | **ND** | **Loose bipolar** | **Partial Co-localized, bipolar** | **Partial Co-localized, bipolar** |
| **CCNA_02255** | **YFP-SpmX** | **Scaffolding** | **ND** | **Diffuse** | **Co-localized, diffuse** | **Co-localized, mostly monopolar** |
| **CCNA_02255** | **SpmX-YFP** | **Scaffolding** | **ND** | **Diffuse** | **Co-localized, diffuse** | **Co-localized, mostly monopolar** |
| **CCNA_02567** | **PleC-mChy** | **Signaling** | **ND** | **Diffuse** | **Co-localized, monopolar** | **No co-localization, diffuse** |
| CCNA_03598 | DivL-mChy | Signaling | ND | Diffuse | No co-localization, diffuse | Co-localized (Holmes et al., 2016) |
| CCNA_03598 | DivL (139-769) -mChy | Signaling | ND | Diffuse | No co-localization, diffuse | Co-localized (Holmes et al., 2016) |
| CCNA_02547 | DivK-mChy | Signaling | PhoB (34%) | Diffuse | No co-localization, diffuse | No co-localization, diffuse |
| CCNA_01132 | CckA-mChy | Signaling | ND | Diffuse | No co-localization, diffuse | Co-localized (Holmes et al., 2016) |
| CCNA_01116 | YFP-DivJ | Signaling | PhoR (31%) | Diffuse | No co-localization, diffuse | No co-localization, diffuse |
| **CCNA_03038** | **mChy-CpaE** | **Signaling** | **ND** | **Diffuse** | **Co-localized, bipolar** | **Partial Co-localized, monopolar** |
| CCNA_02546 | PleD-mChy | Signaling | ND | Diffuse | No co-localization, diffuse | No co-localization, diffuse |
| **CCNA_01918** | **PopA-mChy** | **Signaling** | **ND** | **Diffuse and Spotty** | **Co-localized, bipolar** | **Partial Co-localized, monopolar** |
| **CCNA_02623** | **FtsZ-mChy** | **Cytokinesis** | **FtsZ (52%)** | **Spotty** | **Partial Co-localized, spotty** | **No co-localization, spotty** |
| CCNA_01612 | mChy-MreB | Cytokinesis | MreB (64%) | Diffuse | No co-localization, diffuse | No co-localization, diffuse |
| CCNA_03869 | ParA-mChy | Segregation | ND | Diffuse | No co-localization, diffuse | Co-localized, monopolar |
| CCNA_03868 | CFP-ParB | Segregation | ND | Diffuse | No co-localization, diffuse | Co-localized, monopolar |
| ECD_03570 | IbpA-mChy | Protein aggregation | IbpA (100%) | Spotty | No co-localization, Spotty | No co-localization, Spotty |

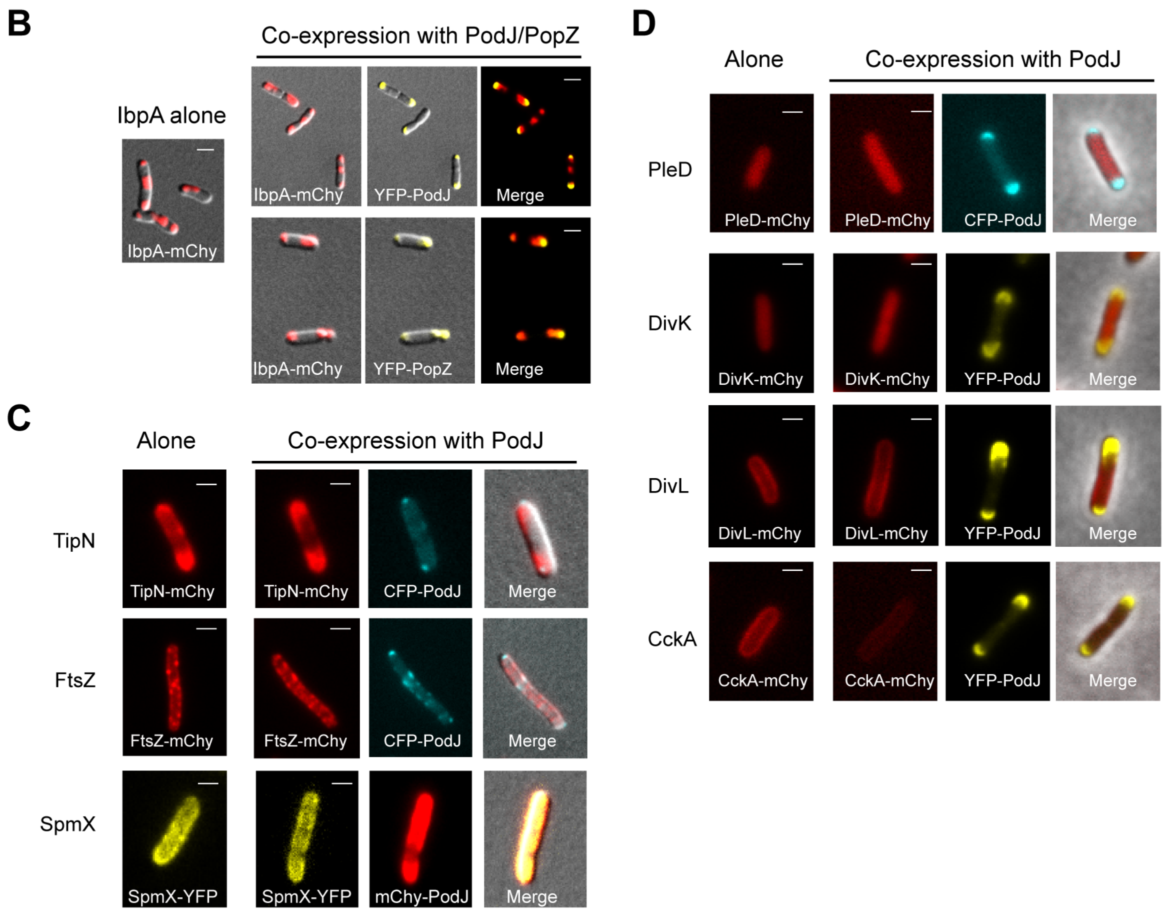


Figure S3: (A) Analysis of PodJ co-expression with potential client proteins in *E. coli*. The proteins in the rows with bold fonts indicate they can interact with PodJ fluorescent protein fusion. (B) Co-expression of YFP-PodJ together with inclusion body protein A (IbpA-mChy) suggests that PodJ does not co-localize with inclusion bodies in *E. coli.* (C) Three potential PodJ protein-protein interaction partners (TipN, FtsZ and SpmX) promoted dispersion of YFP-PodJ when co-expressed in *E. coli.* (D) Co-expression of YFP-PodJ (or CFP-PodJ) together with new cell pole associated proteins (PleD, DivL, DivK, CckA) indicate that these proteins do not co-localize and interact with PodJ in *E. coli.*

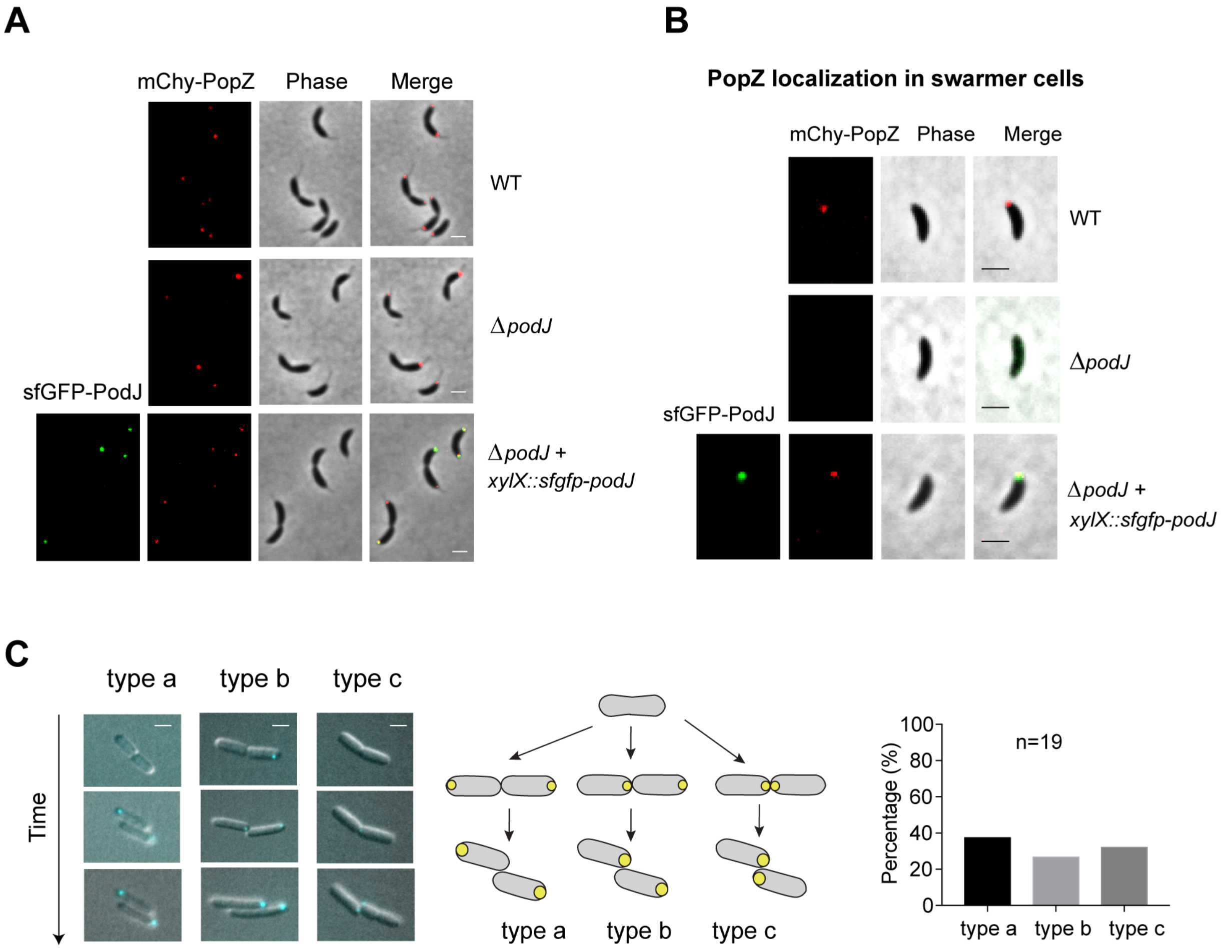


Figure S4: Subcellular localization pattern of mCherry-PopZ in wild-type, Δ*podJ*, and Δ*podJ xylX::sfgfp-podJ* strains in the presence of 0.5 mM vanillate in *C. crescentus* (A) pre-divisional cells and (B) newborn swarmer cells. (C) Time-lapse analyses of mCherry-PopZ in *E. coli* BL21 suggests that PopZ accumulates randomly as a single focus at either the old or new cell poles.

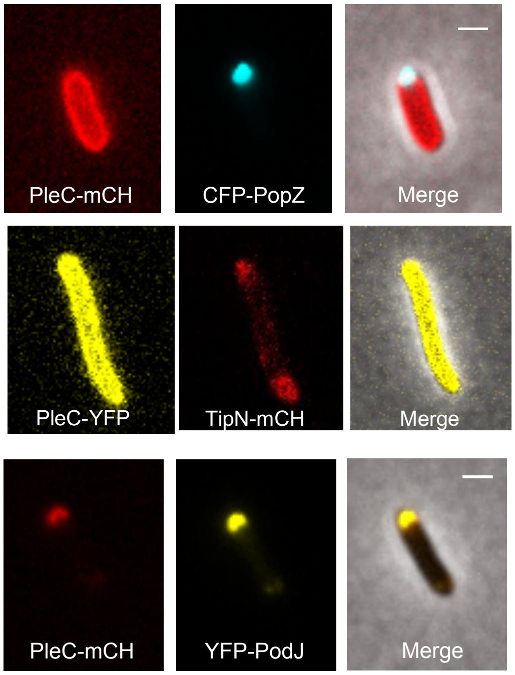


**Figure S5:** Heterologous co-expression of PleC together with 3 new cell-pole associated scaffolds (PopZ, PodJ, and TipN). These assays imply that PleC can be directly recruited to the cell pole by PodJ, while PleC is indirectly associated with the PopZ and TipN scaffold proteins.

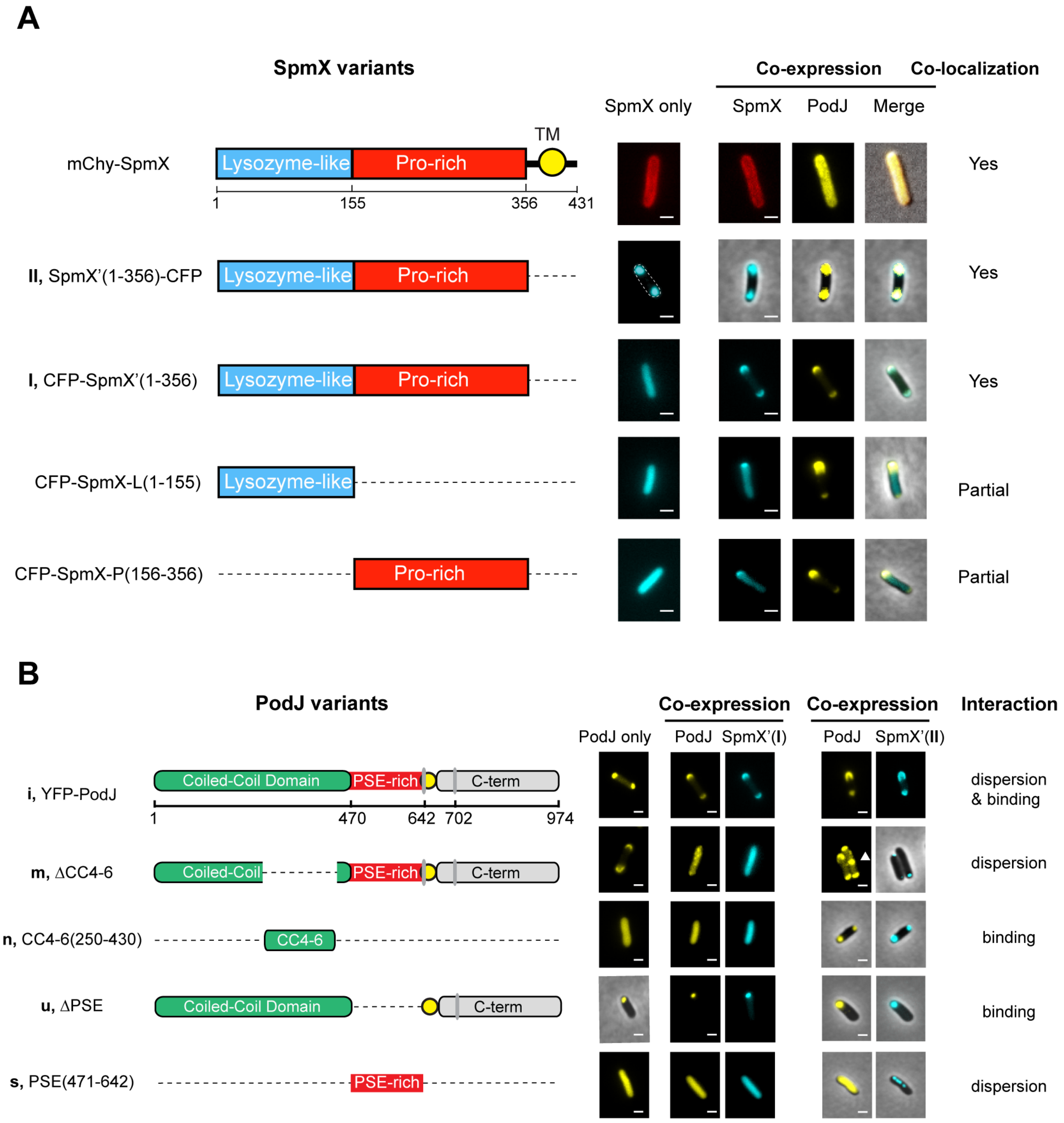


Figure S6: SpmX domain deletion library when expressed alone heterologously in *E. coli,* or co-expressed with PodJ fluorescent protein fusions. Dispersion of YFP-PodJ from the cell pole requires the transmembrane domain of SpmX. The N-terminal fluorescent protein fusion of SpmX(1-356) disrupts its capability to accumulate as a focus suggesting that the N-terminus of SpmX may be involved in self-assembly. The SpmX-PodJ interaction requires both the lysozyme and proline-rich domains of SpmX. (B) Select PodJ domain deletion library variants when expressed alone heterologously in *E. coli* or co-expressed with SpmXΔTM fluorescent protein fusions (please refer the two constructs in Figure S6A: **I**, CFP-SpmXΔTM, and **II**, SpmXΔTM-

CFP). These results suggest that PodJ's PSE and CC4-6 domains are sites of interaction with SpmX.

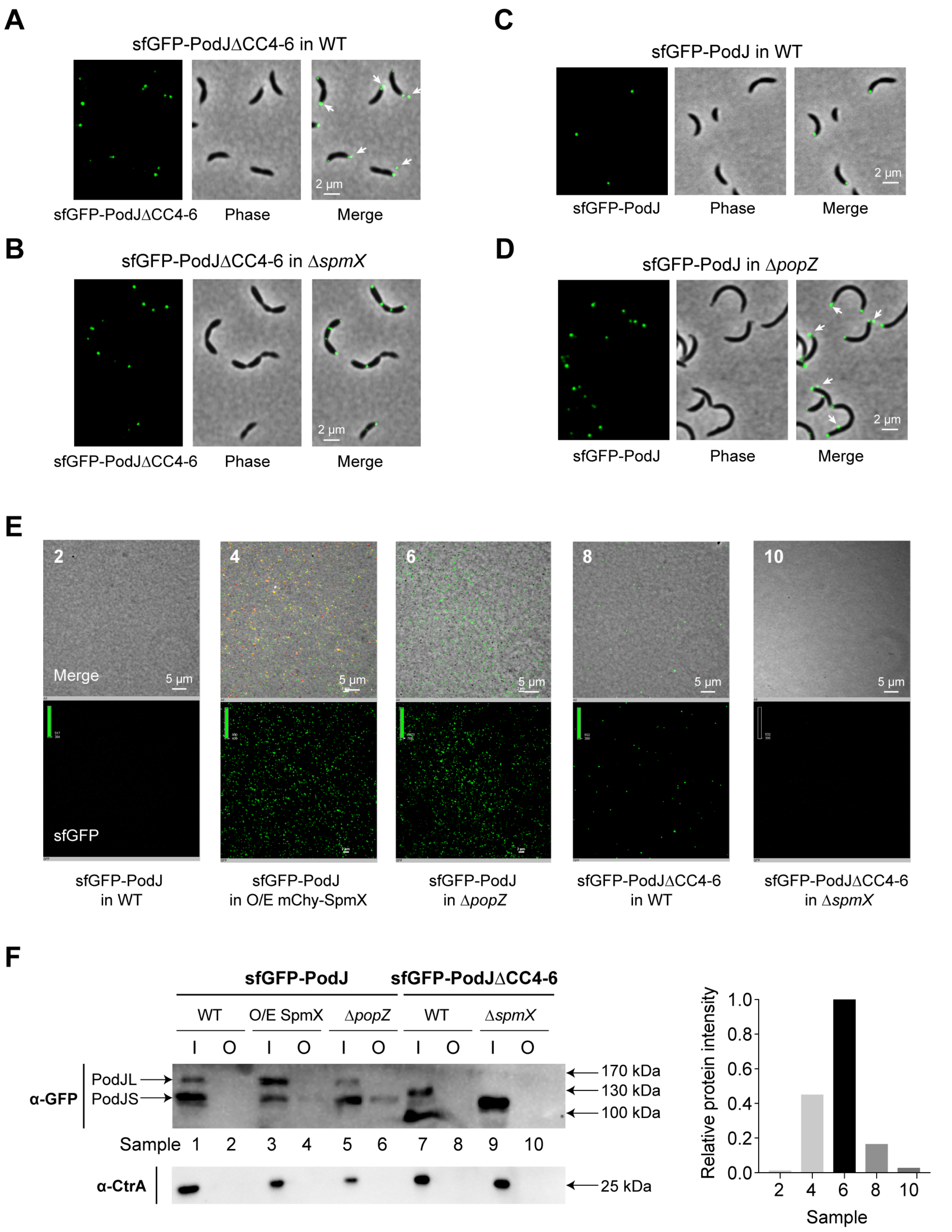


Figure S7. The PopZ-PodJ interaction anchors PodJ in the cytoplasm and prevents PodJ cellular secretion from the old pole. (A) PodJ is specifically secreted from *C. crescentus* strains that disrupt the PodJ-PopZ interaction (PodJΔCC4-6), and (B) the PodJ secretion of C. crescentus requires the SpmX protein. (C and D) Full-length sfGFP-PodJ is also secreted from cells in the PopZ deletion strain (Δ*popZ*). (E) Fractionated media confirms the presence of sfGFP-PodJ foci in the growth media. (F) Western blot of analysis of sfGFP-PodJ and CtrA inside (I) and outside (O) of the cells. The Δ*popZ* strain exhibits the largest amount of extracellular sfGFP-PodJ, while western blot controls of CtrA indicates that the observed PodJ secreted foci is not from cell lysis.

### Table S1: Strains and plasmids

| Name | Relevant genotype/description | Source/reference |
| --- | --- | --- |
| ***C. crescentus* strains** | | |
| WSC0439 | *C. crescentus* NA1000 | Lucy Shapiro |
| WSC1141 | *C. crescentus* NA1000 *parB::CFP-parB, popZ::popZ-mCherry* | (Ptacin et al., 2014) |
| LS3778 | *C. crescentus* NA1000 Δ*podJ42-959* | (Viollier et al., 2002) |
| LS3797 | *C. crescentus* NA1000 Δ*podJ*, Δ*pleC* | (Viollier et al., 2002) |
| LS4367 | *C. crescentus* NA1000 Δ*tipN* | (Huitema et al., 2006) |
| WSC1018 | *C. crescentus* NA1000 Δ*popZ*, *vanA::mCherry-popZ* | (Bowman et al., 2013) |
| WSC1017 | *C. crescentus* NA1000 Δ*spmX* | (Radhakrishnan et al., 2008) |
| WSC1140 | *C. crescentus* NA1000 Δ*pleC* | (Wheeler and Shapiro, 1999) |
| WSC1201 | *C. crescentus* NA1000 Δ*podJ*, *xylX::sfGFP-podJ* | This work |
| WSC1202 | *C. crescentus* NA1000 Δ*podJ*, *xylX::YFP-podJ* | This work |
| WSC1203 | *C. crescentus* NA1000 Δ*podJ*, *podJ::Mng-podJ* | This work |
| WSC1204 | *C. crescentus* NA1000 Δ*podJ*, Δ*pleC*, *xylX::sfGFP-podJ* | This work |
| WSC1205 | *C. crescentus* NA1000 *vanA::mCherry-popZ* | This work |
| WSC1206 | *C. crescentus* NA1000 Δ*podJ*, *vanA::mCherry-popZ* | This work |
| WSC1207 | *C. crescentus* NA1000 Δ*podJ*, *vanA::mCherry-popZ, xylX::sfGFP-podJ* | This work |
| WSC1208 | *C. crescentus* NA1000 Δ*podJ*, *xylX::sfGFP-podJ*Δ*250-430*, *vanA::mCherry-popZ* | This work |
| WSC1209 | *C. crescentus* NA1000/pBVMCS6-Pvan-PleC-mCherry | This work |
| WSC1210 | *C. crescentus* NA1000 Δ*podJ/*pBVMCS6-Pvan-PleC-mCherry | This work |
| WSC1211 | *C. crescentus* NA1000 Δ*podJ*, *xylX::sfGFP-podJ/*pBVMCS6-Pvan-PleC-mCherry | This work |
| WSC1212 | *C. crescentus* NA1000 Δ*tipN/*pBVMCS6-Pvan-PleC-mCherry | This work |
| WSC1213 | *C. crescentus* NA1000 Δ*podJ*, *xylX::sfGFP-podJ/*pBVMCS6-Pvan-mCherry-SpmX | This work |
| WSC1214 | *C. crescentus* NA1000 Δ*podJ*, *xylX::sfGFP-podJ1-635* | This work |
| WSC1215 | *C. crescentus* NA1000 Δ*podJ*, *xylX::sfGFP-podJ*Δ*471-635* | This work |
| WSC1216 | *C. crescentus* NA1000 Δ*podJ*, *xylX::sfGFP-podJ*Δ*250-430* | This work |
| WSC1217 | *C. crescentus* NA1000 Δ*podJ*, *xylX::sfGFP-podJ1-635*, *vanA::mCherry-popZ* | This work |
| WSC1218 | *C. crescentus* NA1000 Δ*podJ*, *xylX::sfGFP-podJ*Δ*471-635*, *vanA::mCherry-popZ* | This work |
| WSC1219 | *C. crescentus* NA1000 Δ*spmX*, *xylX::sfGFP-podJ*Δ*250-430* | This work |
| WSC1220 | *C. crescentus* NA1000 Δ*podJ*, *xylX::sfGFP-podJ1-635*/pBVMCS6-Pvan-PleC-mCherry | This work |
| WSC1221 | *C. crescentus* NA1000 Δ*podJ*, *xylX::sfGFP-podJ*Δ*471-635*/pBVMCS6-Pvan-PleC-mCherry | This work |
| WSC1222 | *C. crescentus* NA1000 Δ*podJ*, *xylX::sfGFP-podJ*Δ*250-430*/pBVMCS6-Pvan-PleC-mCherry | This work |
| WSC1223 | *C. crescentus* NA1000/pBXMCS2-Pxyl-mNeonGreen-PodJ | This work |
| WSC1224 | *C. crescentus* NA1000 Δ*pleC/*pBXMCS2-Pxyl-mNeonGreen-PodJ | This work |
| WSC1225 | *C. crescentus* NA1000 Δ*spmX/*pBXMCS2-Pxyl-mNeonGreen-PodJ | This work |
| WSC1226 | *C. crescentus* NA1000 Δ*popZ*; *vana::mCherry-popZ*/pBXMCS2-Pxyl-mNeonGreen-PodJ | This work |
| WSC1228 | *C. crescentus* NA1000 Δ*tipN, xylX::sfGFP-podJ* | This work |

| | | | |
|---|---|---|---|
| WSC1229 | *C. crescentus* NA1000 Δ*spmX, xylX::sfGFP-podJ* | This work | |
| WSC1230 | *C. crescentus* NA1000 Δ*popZ, vanA::mCherry-popZ, xylX::sfGFP-podJ* | This work | |
| ***E. coli* strains** | | | |
| WSC1123 | BL21 Rosetta, $F^-$ *ompT hsdSB*($Rb^-$ $Mb^-$) *gal dcm* (DE3) Prare (CamR) | Novagen | |
| WSC1119 | DH5α | Novagen | |
| **Plasmids in DH5α** | | **Plasmids** | **Source/reference** |
| WSC1231 | pCDFDuet1-PodJ-YPF | pWZ007 | This work |
| WSC1232 | pCDFDuet1-YFP-PodJ | pWZ012 | This work |
| WSC1233 | pCDFDuet1-mCherry-PodJ | pWZ013 | This work |
| WSC1234 | pBAD-YFP-PodJ | pWZ024 | This work |
| WSC1235 | pBAD-CFP-PodJ | pWZ074 | This work |
| WSC1236 | pCDFDuet1-YFP-PodJ(1-894) | PodJ01 | This work |
| WSC1237 | pCDFDuet1-YFP-PodJ(1-702) | PodJ02 | This work |
| WSC1238 | pCDFDuet1-YFP-PodJ(703-974) | pWZ053 | This work |
| WSC1239 | pCDFDuet1-YFP-PodJ(703-894) | pWZ054 | This work |
| WSC1240 | pCDFDuet1-YFP-PodJ(1-642) | PodJ05 | This work |
| WSC1241 | pCDFDuet1-YFP-PodJ(643-974) | pWZ055 | This work |
| WSC1242 | pCDFDuet1-mCherry-PodJ(1-470) | PodJ07 | This work |
| WSC1243 | pCDFDuet1-mCherry-PodJ(471-642) | PodJ08 | This work |
| WSC1244 | pCDFDuet1-mCherry-PodJ(703-974) | PodJ09 | This work |
| WSC1245 | pCDFDuet1-mCherry-PodJ(703-894) | PodJ10 | This work |
| WSC1246 | pCDFDuet1-mCherry-PodJ(643-974) | PodJ11 | This work |
| WSC1247 | pCDFDuet1-mCherry-PodJ(660-756) | PodJ12 | This work |
| WSC1248 | pCDFDuet1-PodJ-mCherry(703-974) | PodJ13 | This work |
| WSC1249 | pCDFDuet1-mCherry-PodJ(643-756) | PodJ14 | This work |
| WSC1250 | pCDFDuet1-mCherry-PodJ(643-894) | PodJ15 | This work |
| WSC1251 | pCDFDuet1-mCherry-PodJ(471-702) | PodJ16 | This work |
| WSC1252 | pCDFDuet1-YFP-PodJ(1-470, 643-702) | PodJ17 | This work |
| WSC1253 | pCDFDuet1-mCherry-PodJ(643-702) | PodJ18 | This work |
| WSC1254 | pCDFDuet1-YFP-PodJ(128-702) | PodJ19 | This work |
| WSC1255 | pCDFDuet1-YFP-PodJ(170-702) | PodJ20 | This work |
| WSC1256 | pCDFDuet1-YFP-PodJ(250-702) | PodJ21 | This work |
| WSC1257 | pCDFDuet1-YFP-PodJ(430-702) | PodJ22 | This work |
| WSC1258 | pCDFDuet1-YFP-PodJ(1-635) | PodJ23 | This work |
| WSC1259 | pCDFDuet1-mCherry-PodJ(250-430) | pWZ94 | This work |
| WSC1260 | pCDFDuet1-mCherry-PodJ(36-635) | pWZ95 | This work |
| WSC1261 | pCDFDuet1-mCherry-PodJ(36-702) | pWZ118 | This work |
| WSC1262 | pCDFDuet1-YFP-PodJ(42-702) | pWZ202 | This work |
| WSC1263 | pCDFDuet1-YFP-PodJ(1-601) | pWZ203 | This work |

| | | | |
|---|---|---|---|
| WSC1264 | pCDFDuet1-mCherry-PodJ(643-656) | PodJ29 | This work |
| WSC1265 | pCDFDuet1-mCherry-PodJ(657-702) | PodJ30 | This work |
| WSC1266 | pCDFDuet1-YFP-PodJ(1-588, 643-702) | PodJ31 | This work |
| WSC1267 | pCDFDuet1-YFP-PodJ(1-470, popz24-102, 643-702) | PodJ32 | This work |
| WSC1268 | pCDFDuet1-YFP-PodJ(1-470, 589-702) | PodJ33 | This work |
| WSC1269 | pCDFDuet1-YFP-PodJΔ250-430 | PodJ-detal250-430 | This work |
| WSC1273 | pCDFDuet1-mCherry-PopZ | pWZ014 | This work |
| WSC1274 | pBAD-CFP-PopZ | PWZ046 | This work |
| WSC1275 | pCDFDuet1-mCherry-PopZ(1-134) | PopZ1 | This work |
| WSC1276 | pCDFDuet1-mCherry-PopZ(24-134) | PopZ3 | This work |
| WSC1277 | pCDFDuet1-mCherry-PopZ(1-23, 103-134) | PopZ4 | This work |
| WSC1278 | pCDFDuet1-mCherry-PopZ(1-23,103-177) | PopZ5 | This work |
| WSC1279 | pCDFDuet1-mCherry-PopZ(1-102, 135-177) | PopZ6 | This work |
| WSC1280 | pCDFDuet1-mCherry-PopZ(24-177) | PopZ7 | This work |
| WSC1281 | pCDFDuet1-mCherry-PopZ(24-102, 135-177) | PopZ8 | This work |
| WSC1282 | pCDFDuet1-mCherry-PopZ(24-102) | PopZ9 | This work |
| WSC1283 | pCDFDuet1-mCherry-PopZ(1-102) | PopZ2 | This work |
| WSC1284 | pACYCDuet1-YFP-SpmX | pWZ036 | This work |
| WSC1285 | pACYCDuet1-mCherry-SpmX | pWZ037 | This work |
| WSC1286 | pCDFDuet1-CFP-SpmX | PWZ050 | This work |
| WSC1287 | pACYCDuet1-SpmX(1-350)-YFP | pWZ035 | This work |
| WSC1288 | pBAD-SpmX(1-350)-mCherry | pWZ082 | This work |
| WSC1289 | pCDFDuet1-CFP-SpmX(1-356) | pWZ201 | This work |
| WSC1290 | pCDFDuet1-CFP-SpmX(1-155) | pWZ204 | This work |
| WSC1291 | pCDFDuet1-CFP-SpmX(156-356) | pWZ172 | This work |
| WSC1292 | pACYCDuet1-PleC-mCherry | pWZ021 | This work |
| WSC1293 | pBAD-PleC-mCherry | pWZ076 | This work |
| WSC1294 | pBAD-PleC-YFP | pWZ083 | This work |
| WSC1295 | pCDFDuet1-PleC-CFP | pWZ171 | This work |
| WSC1296 | pBAD-PleC(54-842)-mCherry | PleC1 | This work |
| WSC1297 | pBAD-PleC(1-53, 302-842)-mCherry | PleC2 | This work |
| WSC1298 | pBAD-PleC(1-301, 551-842)-mCherry | PleC3 | This work |
| WSC1299 | pBAD-PleC(1-550)-mCherry | PleC4 | This work |
| WSC1300 | pBAD-PleC-pasC-mCherry | pWZ127 | This work |
| WSC1301 | pBAD-PleC-ATG-pasD-mCherry | PWZ135 | This work |
| WSC1302 | pBAD-PleC-pacCD-mCherry | PWZ136 | This work |
| WSC1303 | pBAD-PleCΔpasC | pBAD-plec-ΔpasC | This work |
| WSC1304 | pBAD-PleCΔpasD | pBAD-plec-ΔpasD | This work |
| WSC1305 | pBAD-IbpA-YFP | pWZ085 | This work |

| | | | |
|---|---|---|---|
| WSC1306 | pCDFDuet1-IbpA-mCherry | PWZ047 | This work |
| WSC1307 | pCDFDuet1-IbpA-YFP | PWZ049 | This work |
| WSC1308 | pACYCDuet1-ParA-mCherry | pWZ028 | This work |
| WSC1309 | pACYCDuet1-CFP-ParB | pWZ067 | This work |
| WSC1310 | pBAD-CckA-mCherry | pWZ125 | This work |
| WSC1311 | pBAD-DivK-mCherry | pWZ121 | This work |
| WSC1312 | pBAD-DivL-mCherry | pWZ124 | This work |
| WSC1313 | pACYCDuet1-DivL(139-769)-mCherry | pWZ026 | This work |
| WSC1314 | pACYCDuet1-YFP-DivJ | pWZ058 | This work |
| WSC1315 | pBAD-DivJ-mCherry | pWZ123 | This work |
| WSC1316 | pACYCDuet1-TipN-mCherry | pWZ066 | This work |
| WSC1317 | pACYCDuet1-TipN-CFP | pWZ161 | This work |
| WSC1318 | pACYCDuet1-FtsZ-mCherry | pWZ068 | This work |
| WSC1319 | pACYCDuet1-mCherry-MreB | pWZ070 | This work |
| WSC1320 | pACYCDuet1-mCherry-CpaE | pWZ099 | This work |
| WSC1321 | pACYCDuet1-PopA-mCherry | pWZ101 | This work |
| WSC1322 | pACYCDuet1-PleD-mCherry | pWZ100 | This work |
| WSC1323 | pBAD-YFP-PodJ-PleC-mCherry | pWZ078 | This work |
| WSC1324 | pNPTS138-PodJup-cholor-PodJdw | pWZ166 | This work |
| WSC1325 | pTEV5-CckA | pTEV5-CCKA-cc | This work |
| WSC1326 | pTEV5-DivL | pTEV5-DIVL-cc | This work |
| WSC1327 | pTEV5-PodJ1-635 | pWZ091 | This work |
| WSC1328 | pTEV5-PodJ471-635 | pWZ096 | This work |
| WSC1329 | pTEV5-PodJ250-430 | pWZ098 | This work |
| WSC1330 | pTEV5-Cys-PodJ471-635 | pWZ096-cys | This work |
| WSC1331 | pTEV5-Cys-PodJ250-430 | pWZ098-cys | This work |
| WSC1332 | pTEV5-PleC-pasCD | pWZ137 | This work |

#### Supplemental Experimental Procedures

##### *C. crescentus* Chromosomal DNA Isolation

*C. crescentus* NA1000 chromosomal DNA was isolated as previously described. The NA1000 cells were grown in PYE medium to the logarithmic phase and then collected at OD about 0.4. The cultures were then subjected to lysis using Lysis buffer (10 mM Tris-HCl pH 8.0, 100 mM NaCl, 5 mM EDTA, 0.2 % SDS and 200 µg/ ml Proteinase K). After precipitation of the supernatant with isopropanol, the chromosomal DNA was isolated through centrifugation and resuspended in water.

##### Construction of *C. crescentus* strains

WSC1202: An integrating plasmid (pXYFPN-2-P*xyl*-YFP-PodJ) was constructed and electroporated into the Δ*podJ* strain LS3778, selected for kanamycin resistance. The *podJ* gene was PCR amplified from the genomic DNA of *C. crescentus* NA1000 by using primers WZ0247 and WZ0248. The backbone including the P*xyl* promoter, *yfp* gene and the HRSAT linker was PCR amplified from the plasmid pXYFPN-2 by using primers WZ0010 and WZ0246. The two purified PCR products (100 ng backbone and 200 ng *podJ* fragment) were employed to perform Gibson assembly to generate the plasmid pXYFPN-2-P*xyl*-YFP-PodJ. The selected plasmid with YFP fused at the N terminal of PodJ was verified by sequencing using primers Pxyl-for and M13-for. Correct plasmid (1 µg) was then electroporated and integrated into the *xylX* locus of Δ*podJ* strain and the integration was tested by colony PCR using primers RecXyl-2 and RecUni-1(Thanbichler et al., 2007).

| | |
|---|---|
| WZ0246 | CGGTCTACGCGCGCTAAGCCTTAATTAATATGCATGGTACCTTAAGATCTCG |
| WZ0010 | GGTGGCCGACCGGTGCTTGTACAGCTCGTCCATGCCG |
| WZ0247 | GACGAGCTGTACAAGCACCGGTCGGCCACCATGACG |
| WZ0248 | CCATGCATATTAATTAAGGCTTAGCGCGCGTAGACCGACAGG |
| Pxyl-for | CCCACATGTTAGCGCTACCAAGTGC |
| M13-for | GCCAGGGTTTTCCCAGTCACGA |
| RecXyl-2 | TCTTCCGGCAGGAATTCACTCACGCC |
| RecUni-1 | ATGCCGTTTGTGATGGCTTCCATGTCG |

WSC1201: Similar to the construction of strain WSC1202, an integrating plasmid (pXYFPN-2-P*xyl*-sfGFP-PodJ) was generated from plasmid of pXYFPN-2-P*xyl*-YFP-PodJ and integrated into the *xylX* locus of Δ*podJ* strain LS3778. The *sfgfp* gene was PCR amplified from the plasmid pSR58.6 (Addgene). Primers for Gibson assembly were shown as below. Primers for colony verification were used as in WSC1202 construction.

| | |
|---|---|
| WZ0284_(p0102)_forward | GATGAACTGTACAAACACCGGTCGGCCACCATGACG |
| WZ0285_(p0102)_reverse | GCCTTTACGCATATGGTCGTCTCCCCAAAACTCGAGC |
| WZ0286_(sfGFP)_forward | GTTTTGGGGAGACGACCATATGCGTAAAGGCGAAGAGCTGT |
| WZ0287_(sfGFP)_reverse | GGTGGCCGACCGGTGTTTGTACAGTTCATCCATACCATGCGT |

WSC1204, WSC1228, WSC1229, WSC1230: The integrating plasmid (pXYFPN-2-P*xyl*-sfGFP-PodJ) was electroporated into the *xylX* locus of Δ*podJ*Δ*pleC* (LS3797), Δ*tipN* (LS4367), Δ*spmX* (WSC1017), and Δ*popZ* (WSC1018) strain, generating WSC1204, WSC1228, WSC1229, and WSC1230, respectively. Primers for colony verification were used as in WSC1202 construction.

WSC1203: An integrating plasmid (pXYFPN-2-P*podJ*-mNG-PodJ) was constructed and electroporated into the Δ*podJ* strain LS3778, selected for kanamycin resistance. The backbone including the full length *podJ* gene and the HRSAT linker but without P*xyl* promoter and *yfp* gene was PCR amplified from the plasmid (pXYFPN-2-P*xyl*-YFP-PodJ) by using primers WZ0170 and WZ0263. The promoter of *podJ* (500 bp upstream of the ATG) was PCR amplified from the genomic DNA of *C. crescentus* by using primers WZ0264 and WZ0265. The *mNeonGreen* gene was PCR amplified from the plasmid pNCS-mNeonGreen (Allele Biotechnology) by using primers WZ0266 and WZ0267. The purified backbone and inserts (100 ng backbone and 200 ng insert fragments) were employed to perform Gibson assembly to generate the plasmid pXYFPN-2-P*podJ*-mNG-PodJ. The verified correct plasmid (1 μg) was then electroporated into the Δ*podJ* strain and the integration was screened using colony PCR with primers PODJ-KO-C-F and PODJ-KO-C-R flanking the native *podJ* gene.

| | |
|---|---|
| WZ0170_(pbad-podj)_forward | GGACGAGCTGTACAAGCACCGGTCGGCCACCATGAC |
| WZ0263_(pxyfpn-2-podj)_reverse | CCTGATCCTCGCCGATCCGCTGCCTGGTGCTGGACC |
| WZ0264_(Ppodj)_forward | GCACCAGGCAGCGGATCGGCGAGGATCAGGAACAGG |
| WZ0265_(Ppodj)_reverse | GCCCTTGCTCACCATGCGAATCGATCTCCCCGCACC |
| WZ0266_(mNeonGreen)_forward | GGGAGATCGATTCGCATGGTGAGCAAGGGCGAGGAGGA |
| WZ0267_(mNeonGreen)_reverse | GGTGGCCGACCGGTGCTTGTACAGCTCGTCCATGCCCA |
| PODJ-KO-C-F | CTTCTAGGCCTGCGACAATC |
| PODJ-KO-C-R | TGGAGTTTATCCCGAACAGG |

WSC1214, WSC1215, WSC1216: Three integrating plasmids were constructed, respectively, *i.e.*, pXYFPN-2-P*xyl*-sfGFP-PodJ1-635, pXYFPN-2-P*xyl*-sfGFP-PodJΔ471-635, and pXYFPN-2-P*xyl*-sfGFP-PodJΔ250-430. They used the same template plasmid pXYFPN-2-P*xyl*-sfGFP-PodJ but used different overlapping primers (see below). The purified PCR products (200 ng) were transformed directly into DH5α, selecting for kanamycin resistance. The selected correct plasmids were verified by sequencing using primer podj-422-F, podj-1647-R, podj-1520-F, or podj-2522-R. Correct plasmids (1 μg) were then electroporated and integrated into the *xylX* locus of Δ*podJ* strain, generating WSC1214 (PodJ1-635), WSC1215 (PodJΔ471-635), and WSC1216 (PodJΔ250-430), respectively. The integration was tested by colony PCR using primers RecXyl-2 and RecUni-1(Thanbichler et al., 2007).

| | |
|---|---|
| **WZH566-pxyfpn-2-Pxyl-sfGFP-PodJ1-635-F** | **GGCGCGCTTGTAAGCCTTAATTAATATGCA** |
| **WZH567-pxyfpn-2-Pxyl-sfGFP-PodJ1-635-R** | TTAAGGCTTACAAGCGCGCCTTCGACTTCT |
| **WZH568-pxyfpn-2-Pxyl-sfGFP-PodJΔ471-635-F** | AGCTGCGCCGGGCGCGACCGTGACGACGGC |
| **WZH569-pxyfpn-2-Pxyl-sfGFP-PodJΔ471-635-R** | CGGTCGCGCCCGGCGCAGCTTCCAGCTTCC |
| **WZH570-pxyfpn-2-Pxyl-sfGFP-PodJΔ250-430-F** | GGATCAGCGCCAGGAACTGGTCGACCGCAT |
| **WZH571-pxyfpn-2-Pxyl-sfGFP-PodJΔ250-430-R** | CCAGTTCCTGGCGCTGATCCAGGGCCTGGA |
| **WZ142-podj-422-F** | ACGAACTGAAGACCGAGCAG |
| **WZ143-podj-1647-R** | GATCGCGTAGTCATCGTGTG |
| **WZ144-podj-1520-F** | TCAGCACGTCTGAGGATGAG |
| **WZ145-podj-2522-R** | TCGTAGAGCTGAGCCAGGTT |

WSC1219: The integrating plasmid (pXYFPN-2-P*xyl*-sfGFP-PodJΔ250-430) was electroporated into the *xylX* locus of Δ*spmX* strain WSC1017. Primers for colony verification were used as in WSC1216 construction.

WSC1205, WSC1206, WSC1207, WSC1208, WSC1217, WSC1218: An integrating plasmid (pVCHYN-6-P*van*-mCherry-PopZ) was constructed, selected for chloramphenicol resistance. The *mCherry-popZ* gene was PCR amplified from the plasmid WSC1273 (pCDFDuet1-mCherry-PopZ) by using primers WZ0330 and WZ0331. The backbone including the P*van* promoter and the HRSAT linker was PCR amplified from the plasmid pVCHYN-6 by using primers WZ0328 and WZ0329. The two purified PCR products (100 ng backbone and 200 ng *mCherry-popZ* fragment) were employed to perform Gibson assembly to generate the plasmid pVCHYN-6-P*van*-mCherry-PopZ. The selected plasmid with mCherry fused at the N terminal of PopZ was verified by sequencing using primers Pvan-for and M13-for (Thanbichler et al., 2007). Sequence verified plasmid (1 μg) was then electroporated and integrated into the *van* locus of the wild-type, Δ*podJ* (LS3778), WSC1201, WSC1214, WSC1215, and WSC1216 strain, generating WSC1205, WSC1206, WSC1207, WSC1217, WSC1208, and WSC1218 strain, respectively. The integration was tested by colony PCR using primers RecVan-2 and RecUni-1 (Thanbichler et al., 2007).

| | |
|---|---|
| WZ0328_(pvan)_forward | GGACGCGGCGCCTAATCTAGAGCGGCCATTCACTGGCC |
| WZ0329_(pvan)_reverse | GCCCTTGCTCACCATATGCGTTTCCTCGCATCGTGGTTCG |
| WZ0330_(mch-popz)_forward | TGCGAGGAAACGCATATGGTGAGCAAGGGCGAGGAGG |
| WZ0331_(mch-popz)_reverse | AATGGCCGCTCTAGATTAGGCGCCGCGTCCCCGAG |
| Pvan-for | GACGTCCGTTTGATTACGATCAAGATTGG |
| M13-for | GCCAGGGTTTTCCCAGTCACGA |
| RecVan-2 | CAGCCTTGGCCACGGTTTCGGTACC |
| RecUni-1 | ATGCCGTTTGTGATGGCTTCCATGTCG |

WSC1209, WSC1210, WSC1211, WSC1212, WSC1220, WSC1221, WSC1222: A replicating plasmid (pBVMCS6-P*van*-PleC-mCherry) was constructed, selecting for chloramphenicol resistance. The *plec-mCherry* gene was PCR amplified from the plasmid WSC1293 (pBAD-PleC-mCherry) by using primers WZ0326 and WZ0327. The backbone including the P*van* promoter and the HRSAT linker was PCR amplified from the plasmid pBVMCS-6 by using primers WZ0324 and WZ0325. The two purified PCR products (100 ng backbone and 200 ng *plec-mCherry* fragment) were employed to perform Gibson assembly to generate the plasmid pBVMCS6-P*van*-PleC-mCherry. The selected plasmid with mCherry fused at the C terminal of PleC was verified by sequencing using primers Pvan-for and M13-for. Correct plasmid (100 ng) was then electroporated into the wild-type, Δ*podJ* (LS3778), WSC1201, Δ*tipN* (LS4367), WSC1214, WSC1215, and WSC1216 strain, generating WSC1209, WSC1210, WSC1211, WSC1212, WSC1220, WSC1221, and WSC1222 strain, respectively.

| | |
|---|---|
| WZ0324_(pbvmcs-6)_forward | CGAGCTGTACAAGTAAAAAACGGGCCCCCCCTCGAGG |
| WZ0325_(pbvmcs-6)_reverse | CCCGTGTCTGCCCATCGTTTCCTCGCATCGTGGTTCGG |
| WZ0326_(plec-mch)_forward | CGATGCGAGGAAACGATGGGCAGACACGGGGGGCC |
| WZ0327_(plec-mch)_reverse | GGGGGGCCCGTTTTTTACTTGTACAGCTCGTCCATGCCG |

WSC1213: A replicating plasmid (pBVMCS6-Pvan-mCherry-SpmX) was constructed, selected for chloramphenicol resistance. The *mCherry-spmX* gene was PCR amplified from the plasmid WSC1285 (pACYC-mCherry-SpmX) by using primers WZ0261 and WZ0262. The backbone including the P*van* promoter and the HRSAT linker was PCR amplified from the plasmid pBVMCS-6 by using primers WZ0259 and WZ0260. The two purified PCR products (100 ng backbone and 200 ng *mCherry-spmX* fragment) were employed to perform Gibson assembly to generate the plasmid pBVMCS6-Pvan-mCherry-SpmX. The selected plasmid with mCherry fused at the N terminal of SpmX was verified by sequencing using primers Pvan-for and M13-for. Correct plasmid (100 ng) was then electroporated into the WSC1201 cells selecting for chloramphenicol resistance and tested by colony PCR using primers Pvan-for and M13-for (Thanbichler et al., 2007).

| | |
|---|---|
| WZ0259_(pbvmcs-6)_forward | GAGCGACGAAGAGTAGAAAACGGGCCCCCCCTCGAGG |
| WZ0260_(pbvmcs-6)_reverse | GCCCTTGCTCACCATCGTTTCCTCGCATCGTGGTTCGG |
| WZ0261_(mcherry-spmx)_forward | CGATGCGAGGAAACGATGGTGAGCAAGGGCGAGGAGG |
| WZ0262_(mcherry-spmx)_reverse | GGGGGGGCCCGTTTTCTACTCTTCGTCGCTCACATCGGGG |

WSC1223, WSC1224, WSC1225, WSC1226: A replicating plasmid (pBXMCS-2-P*xyl*-mNG-PodJ) was constructed and selected for kanamycin resistance. The backbone including the P*xyl* promoter was PCR amplified from the plasmid pBXMCS-2 by using primers WZ0277 and WZ0278. The *mNeonGreen-podJ* gene was PCR amplified from the plasmid pXYFPN-2-P*podJ*-mNG-PodJ (WSC1203) by using primers WZ0279 and WZ0280. The purified backbone and insert (100 ng backbone and 200 ng *mNeonGreen-podJ* fragment) were employed to perform Gibson assembly to generate the plasmid pBXMCS-2-P*xyl*-mNG-PodJ. The verified correct plasmid (100 ng) was then electroporated into the wild-type, Δ*pleC* (WSC1140), Δ*spmX* (WSC1017), and Δ*popZ* (WSC1018) strain, generating WSC1223, WSC1224, WSC1225, and WSC1226, respectively.

WSC1324: To create an in-frame deletion of *podJ* on the chromosome of *C. crescentus* WSC1141, we constructed a two-step deletion plasmid pNPTS138-PodJup-chlor-PodJdw. We first PCR amplified the upstream (PodJup, 1000 bp) and the downstream (PodJdw, 1000 bp) of *podJ* gene, as well as the antibiotic resistance gene *cat.* These three fragments were purified and inserted into the backbone pNPTS138 via Gibson assembly. The selected plasmid was tested by PCR using primers pnpts138-C-F and pnpts138-C-R and then verified by sequencing. The verified correct plasmid (1 μg) was electroporated into *C. crescentus* WSC1141 and selected for single cross-over colony first ($Kan^R$, $Chlor^R$). The verified colonies were grown overnight in PYE and plated on PYE plate with 3% sucrose and 1 μg/ml chloramphenicol to lose the intervening plasmid. Then, colonies were negatively selected by streaking on PYE plates with kanamycin resistance (25 μg/ml), and the double cross-over colonies ($Kan^S$, $Chlor^R$) were further verified using primer PODJ-KO-C-F and PODJ-KO-C-R.

| | |
|---|---|
| WZ0410_(pnpts138)_forward | GTAGAGGACCGCGTCGTCTAGTCAAGGCCTTAAGTGAGTCG |
| WZ0411_(pnpts138)_reverse | GCTCGTCCATCATGGTCTTCAATTGCACGGGCCCCAC |
| WZ0412_(podj-up)_forward | GCCCGTGCAATTGAAGACCATGATGGACGAGCTGAAGTC |
| WZ0413_(podj-up)_reverse | CCAAAATCCCTTAACGCGAATCGATCTCCCCGCACC |
| WZ0414_(chlor)_forward | GGGAGATCGATTCGCGTTAAGGGATTTTGGTCATCGAACCCCA |
| WZ0415_(chlor)_reverse | GAGGTCGCGAGGCGTTTACGCCCCGCCCTGCCACT |
| WZ0416_(podj-down)_forward | CAGGGCGGGGCGTAAACGCCTCGCGACCTCGCGCT |
| WZ0417_(podj-down)_reverse | CTTAAGGCCTTGACTAGACGACGCGGTCCTCTACAAGGAGAGA |
| pnpts138-C-F | TGCTTCCGGCTCGTATGTTG |
| pnpts138-C-R | GTAATACGACTCACTTAAGG |
| PODJ-KO-C-F | CTTCTAGGCCTGCGACAATC |
| PODJ-KO-C-R | TGGAGTTTATCCCGAACAGG |

All the plasmids used in this study were built in *E. coli* DH5α through Gibson assembly approach. We take WSC1232 as an example to show how to construct these plasmids here. To generate the plasmid pCDFDuet1-YFP-PodJ in WSC1232 strain, we linearized the backbone pCDFDuet1 at *Nde* I by restriction digestion and PCR amplified the two inserts *yfp* and *podJ* gene from plasmid pXYFPC-6 and from genomic DNA, respectively. In this case, the HRSAT linker between YFP and PodJ was designed to be embed in the forward primer for *podJ* gene (WZ0011). The purified backbone and inserts (100 ng backbone and 200 ng insert fragments) were employed to perform Gibson assembly (One-step isothermal reaction). The selected plasmid was tested by colony PCR using primers T7_for and T7_ter, and confirmed by sequencing. The verified correct plasmid (100 ng) was then transformed into BL21 for imaging experiments or protein expression.

| | |
|---|---|
| WZ0025 | CTTAGTATATTAGTTAAGTATAAGAAGGAGATATAATGGTGAGCAAGGGCGAGGAGC |
| WZ0010 | GGTGGCCGACCGGTGCTTGTACAGCTCGTCCATGCCG |
| WZ0011 | GACGAGCTGTACAAGCACCGGTCGGCCACCATGACGGCGGCTTCGCCATGG |
| WZ0026 | GTGGCCGGCCGATATCCAATTGAGATCTGCTTAGCGCGCGTAGACCGACAGG |
| T7_for | GGATCTCGACGCTCTCCCT |
| T7_ter | GCTAGTTATTGCTCAGCGG |

#### All the other primers used to build plasmids in Table S1 are listed as below:

| Primer name | Primer sequence | Plasmids |
|---|---|---|
| **WZ0013_(podj)_forward** | CTTAGTATATTAGTTAAGTATAAGAAGGAGATATAATGACGGCGGCTTCGCCATGG | pWZ007 |
| **WZ0004_(podJ)_reverse** | GGTGGCCGACCGGTGGCGCGCGTAGACCGACAGG | |
| **WZ0014_(hrsat-yfp)_forward** | TCGGTCTACGCGCGCCACCGGTCGGCCACCATGG | |
| **WZ0015_(hrsat-yfp)_reverse** | GTGGCCGGCCGATATCCAATTGAGATCTGCTTACTTGTACAGCTCGTCCATGCCG | |
| **WZ0027_(pxchyc-6)_forward** | CTTAGTATATTAGTTAAGTATAAGAAGGAGATATAATGGTGAGCAAGGGCGAGGAGG | pWZ013 |
| **WZ0010_(PCDF-YFP)_reverse** | GGTGGCCGACCGGTGCTTGTACAGCTCGTCCATGCCG | |
| **WZ0011_(HRSAT)_(PODJ)_forward** | GACGAGCTGTACAAGCACCGGTCGGCCACCATGACGGCGGCTTCGCCATGG | |
| **WZ0026_(PodJ)_reverse** | GTGGCCGGCCGATATCCAATTGAGATCTGCTTAGCGCGCGTAGACCGACAGG | |
| **WZ0028_(mcherry)_forward** | CAGCAGCCATCACCATCATCACCACAGCCAATGGTGAGCAAGGGCGAGGAGG | pWZ014 |
| **WZ0029_(mcherry)_reverse** | GAGACTGATCGGACATGGTGGCCGACCGGTGCTTGTACAGC | |
| **WZ0030_(popz)_forward** | CACCGGTCGGCCACCATGTCCGATCAGTCTCAAGAACCTACAATGG | |
| **WZ0031_(popz)_reverse** | GTCGACCTGCAGGCGCGCCGAGCTCGAATTCTTAGGCGCCGCGTCCCCG | |
| **WZ0044_(Plec)_forward** | CAGCAGCCATCACCATCATCACCACAGCCAATGGGCAGACACGGGGGGC | pWZ021 |
| **WZ0045_(Plec)_reverse** | GGTGGCCGACCGGTGGGCCGCCACGAAGTCGCG | |
| **WZ0046_(mcherry)_forward** | GACTTCGTGGCGGCCCACCGGTCGGCCACCATGG | |
| **WZ0024_(mcherry)_reverse** | GTCGACCTGCAGGCGCGCCGAGCTCGAATTCTTACTTGTACAGCTCGTCCATGCCGC | |
| **WZ0050_(PBAD)_forward** | GGTCTACGCGCGCTAAGAAGCTTGGCTGTTTTGGCGG | pWZ024 |
| **WZ0051_(PBAD)_reverse** | CGCCCTTGCTCACCATTTAATTCCTCCTGTTAGCCCAAAAA | |
| **WZ0052_(YFP-PODJ)_forward** | AACAGGAGGAATTAAATGGTGAGCAAGGGCGAGGAGC | |
| **WZ0053_(YFP-PODJ)_reverse** | AAACAGCCAAGCTTCTTAGCGCGCGTAGACCGACAGG | |
| **WZ0056_(divL139-769)_forward** | CAGCAGCCATCACCATCATCACCACAGCCAATGGCGGGCGCCCTGGCCT | pWZ026 |
| **WZ0017_(divl-cc-26-769)_reverse** | GGTGGCCGACCGGTGGAAGCCGAGTTCGGGCTGCA | |
| **WZ0057_(mcherry)_forward** | CCCGAACTCGGCTTCCACCGGTCGGCCACCATGG | |
| **WZ0058_(mcherry)_reverse** | CTTTCTGTTCGACTTAAGCATTATGCGGCCGCTTACTTGTACAGCTCGTCCATGCCGC | |
| **WZ0060_(parA)_forward** | CAGCAGCCATCACCATCATCACCACAGCCAGTGTCCGCTAATCCTCTCCGCGTTCT | pWZ028 |
| **WZ0061_(parA)_reverse** | GGTGGCCGACCGGTGGGCGGCCTTGGCCTGGCGAT | |
| **WZ0062_(mcherry)_forward** | CAGGCCAAGGCCGCCCACCGGTCGGCCACCATGG | |
| **WZ0058_(mcherry)_reverse** | CTTTCTGTTCGACTTAAGCATTATGCGGCCGCTTACTTGTACAGCTCGTCCATGCCGC | |
| **WZ0072_(spmx-1-350)_forward** | AGCAGCCATCACCATCATCACCACAGCCAGATGAAACCGCGTCATCAGGTCTCCC | pWZ035 |
| **WZ0073_(spmx-1-350)_reverse** | GGTGGCCGACCGGTGGTCCATCACGCCGACGGTTGC | |
| **WZ0074_(yfp)_forward** | GTCGGCGTGATGGACCACCGGTCGGCCACCATGG | |
| **WZ0059_(yfp)_reverse** | CTTTCTGTTCGACTTAAGCATTATGCGGCCGCTTACTTGTACAGCTCGTCCATGCCG | |
| **WZ0075_(yfp)_forward** | AGCAGCCATCACCATCATCACCACAGCCAGATGGTGAGCAAGGGCGAGGAGC | pWZ036 |
| **WZ0010_(PCDF-YFP)_reverse** | GGTGGCCGACCGGTGCTTGTACAGCTCGTCCATGCCG | |
| **WZ0076_(hrsat)_(spmx)_forward** | GACGAGCTGTACAAGCACCGGTCGGCCACCATGAAACCGCGTCATCAGGTCTCCC | |
| **WZ0077_(spmx)_reverse** | CTTTCTGTTCGACTTAAGCATTATGCGGCCGCCTACTCTTCGTCGCTCACATCGGGG | |
| **WZ0078_(mcherry)_forward** | AGCAGCCATCACCATCATCACCACAGCCAGATGGTGAGCAAGGGCGAGGAGG | pWZ037 |
| **WZ0010_(PCDF-YFP)_reverse** | GGTGGCCGACCGGTGCTTGTACAGCTCGTCCATGCCG | |
| **WZ0076_(hrsat)_(spmx)_forward** | GACGAGCTGTACAAGCACCGGTCGGCCACCATGAAACCGCGTCATCAGGTCTCCC | |
| **WZ0077_(spmx)_reverse** | CTTTCTGTTCGACTTAAGCATTATGCGGCCGCCTACTCTTCGTCGCTCACATCGGGG | |
| **WZ0079_(spmx)_forward** | CTTAGTATATTAGTTAAGTATAAGAAGGAGATATAATGAAACCGCGTCATCAGGTCTCCC | pWZ038 |
| **WZ0080_(spmx)_reverse** | GCTACCACTGCCACCCTCTTCGTCGCTCACATCGGGGC | |
| **WZ0081_(ggsgs)_(yfp)_forward** | GTGAGCGACGAAGAGGGTGGCAGTGGTAGCATGGTGAGCAAGGGCGAGGAGC | |
| **WZ0015_(hrsat-yfp)_reverse** | GTGGCCGGCCGATATCCAATTGAGATCTGCTTACTTGTACAGCTCGTCCATGCCG | |

| | | |
|---|---|---|
| **WZ0094_(ftsz)_forward** | CAGCAGCCATCACCATCATCACCACAGCCAATGGCTATTTCTCTTTCCGCGCCGC | pWZ043 |
| **WZ0095_(ftsz)_reverse** | GGTGGCCGACCGGTGGTTGGCCAGGCGGCGCAGG | |
| **WZ0096_(mcherry)_forward** | CGCCGCCTGGCCAACCACCGGTCGGCCACCATGG | |
| **WZ0058_(mcherry)_reverse** | CTTTCTGTTCGACTTAAGCATTATGCGGCCGCTTACTTGTACAGCTCGTCCATGCCGC | |
| **WZ0099_(pbad)_forward** | GGACGCGGCGCCTAAGAAGCTTGGCTGTTTTGGCGG | pWZ046 |
| **WZ0051_(PBAD)_reverse** | CGCCCTTGCTCACCATTTAATTCCTCCTGTTAGCCCAAAAA | |
| **WZ0052_(YFP-PODJ)_forward** | AACAGGAGGAATTAAATGGTGAGCAAGGGCGAGGAGC | |
| **WZ0103_(cfp)_(hrsat)_reverse** | GACTGATCGGACATGGTGGCCGACCGGTGCTTGTACAGCTCGTCCATGCCG | |
| **WZ0104_(popz)_forward** | GCACCGGTCGGCCACCATGTCCGATCAGTCTCAAGAACCTACAATGG | |
| **WZ0105_(popz)_reverse** | CAAAACAGCCAAGCTTCTTAGGCGCCGCGTCCCCGAG | |
| **WZ0106_(ibpA)_forward** | CTTAGTATATTAGTTAAGTATAAGAAGGAGATATAATGCGTAACTTTGATTTATCCCCGC | pWZ047 |
| **WZ0107_(ibpA)_reverse** | GGTGGCCGACCGGTGGTTGATTTCGATACGGCGCGG | |
| **WZ0108_(mcherry)_forward** | CGTATCGAAATCAACCACCGGTCGGCCACCATGGTGAGC | |
| **WZ0015_(hrsat-yfp)_reverse** | GTGGCCGGCCGATATCCAATTGAGATCTGCTTACTTGTACAGCTCGTCCATGCCG | |
| **WZ0106_(ibpA)_forward** | CTTAGTATATTAGTTAAGTATAAGAAGGAGATATAATGCGTAACTTTGATTTATCCCCGC | pWZ049 |
| **WZ0107_(ibpA)_reverse** | GGTGGCCGACCGGTGGTTGATTTCGATACGGCGCGG | |
| **WZ0109_(yfp)_forward** | CGTATCGAAATCAACCACCGGTCGGCCACCATGGTG | |
| **WZ0015_(hrsat-yfp)_reverse** | GTGGCCGGCCGATATCCAATTGAGATCTGCTTACTTGTACAGCTCGTCCATGCCG | |
| **WZ0113_(pWZ013)_forward** | GGACGAGCTGTACAAGTAAGCAGATCTCAATTGGATATCGGCCGGC | pWZ052 |
| **WZ0114_(pWZ013)_reverse** | GGTGGCCGACCGGTGGCGCGCGTAGACCGACAGGC | |
| **WZ0115_(yfp)_forward** | TCGGTCTACGCGCGCCACCGGTCGGCCACCATGGTG | |
| **WZ0116_(yfp)_reverse** | TCCAATTGAGATCTGCTTACTTGTACAGCTCGTCCATGCCGAGA | |
| **WZ182-podj01-f** | CTTGGCCTTCTAAGCAGATCTCAATTGGAT | podj01 |
| **WZ183-podj01-r** | GATCTGCTTAGAAGGCCAAGGCCGAGCGGT | |
| **WZ184-podj02-f** | GCGCGCCGCCTAAGCAGATCTCAATTGGAT | podj02 |
| **WZ185-podj02-r** | GATCTGCTTAGGCGGCGCGCGGCGCGCCGG | |
| **WZ186-podj05-f** | GACGACGGCCTAAGCAGATCTCAATTGGAT | podj05 |
| **WZ187-podj05-r** | GATCTGCTTAGGCCGTCGTCACGGTCGCGC | |
| **WZ0117_(pcdf-yfp)_forward** | GTCTACGCGCGCTAAGCAGATCTCAATTGGATATCGGCCGGC | pWZ053/ podj03 |
| **WZ0118_(pcdf-yfp)_reverse** | GTCGTCAGCGCGACGGTGGCCGACCGGTGCTTGTA | |
| **WZ0119_(podj703-974)_forward** | GCACCGGTCGGCCACCGTCGCGCTGACGACGGGCAAGG | |
| **WZ0120_(podj703-974)_reverse** | CCAATTGAGATCTGCTTAGCGCGCGTAGACCGACAGG | |
| **WZ0121_(pcdf-yfp)_forward** | GGCCTTGGCCTTCTAAGCAGATCTCAATTGGATATCGGCCGGC | pWZ054/ podj04 |
| **WZ0122_(pcdf-yfp)_reverse** | CGTCGTCAGCGCGACGGTGGCCGACCGGTGCTTGTA | |
| **WZ0123_(podj703-894)_forward** | CACCGGTCGGCCACCGTCGCGCTGACGACGGGCAA | |
| **WZ0124_(podj703-894)_(taa)_reverse** | CCAATTGAGATCTGCTTAGAAGGCCAAGGCCGAGCGGT | |
| **WZ0125_(pcdf-yfp)_forward** | CGGTCTACGCGCGCTAAGCAGATCTCAATTGGATATCGGCCGGC | pWZ055/ podj06 |
| **WZ0126_(pcdf-yfp)_reverse** | GCGAAGACAACGAGGGTGGCCGACCGGTGCTTGTA | |
| **WZ0127_(podj643-974)_forward** | GCACCGGTCGGCCACCCTCGTTGTCTTCGCCGCCGC | |
| **WZ0120_(podj703-974)_reverse** | CCAATTGAGATCTGCTTAGCGCGCGTAGACCGACAGG | |
| **WZ220-Podj07-f** | AGCTGCGCCGTAAGCAGATCTCAATTGGAT | podj07 |
| **WZ221-Podj07-r** | GATCTGCTTACGGCGCAGCTTCCAGCTTCC | |
| **WZ0135_(pcdf-mcherry)_forward** | ACCGTGACGACGGCCTAAGCAGATCTCAATTGGATATCGGCCGGC | pWZ059/ podj08 |
| **WZ0136_(pcdf-mcherry)_reverse** | CGGCGACGGGGTGGCCGACCGGTGCTTGTA | |

| | | |
|---|---|---|
| **WZ0137_(podj08)_forward** | TACAAGCACCGGTCGGCCACCCCGTCGCCGCCGCCCGCGCAGG | |
| **WZ0138_(podj08)_reverse** | AATTGAGATCTGCTTAGGCCGTCGTCACGGTCGCGC | |
| **WZ0141_(pcdf-mcherry)_forward** | CCTGTCGGTCTACGCGCGCTAAGCAGATCTCAATTGGATATCGGCCGGC | pWZ061/ podj09 |
| **WZ0122_(pcdf-yfp)_reverse** | CGTCGTCAGCGCGACGGTGGCCGACCGGTGCTTGTA | |
| **WZ0123_(podj703-894)_forward** | CACCGGTCGGCCACCGTCGCGCTGACGACGGGCAA | |
| **WZ0142_(podj)_reverse** | CAATTGAGATCTGCTTAGCGCGCGTAGACCGACAGGC | |
| **WZ0143_(pcdf-mcherry)_forward** | GCTCGGCCTTGGCCTTCTAAGCAGATCTCAATTGGATATCGGCCGGC | pWZ062/ podj10 |
| **WZ0122_(pcdf-yfp)_reverse** | CGTCGTCAGCGCGACGGTGGCCGACCGGTGCTTGTA | |
| **WZ0123_(podj703-894)_forward** | CACCGGTCGGCCACCGTCGCGCTGACGACGGGCAA | |
| **WZ0144_(podj)_reverse** | CAATTGAGATCTGCTTAGAAGGCCAAGGCCGAGCGGT | |
| **WZ0141_(pcdf-mcherry)_forward** | CCTGTCGGTCTACGCGCGCTAAGCAGATCTCAATTGGATATCGGCCGGC | pWZ063/ podj11 |
| **WZ0145_(pcdf-mcherry)_reverse** | GGCGAAGACAACGAGGGTGGCCGACCGGTGCTTGTA | |
| **WZ0146_(podj)_forward** | CACCGGTCGGCCACCCTCGTTGTCTTCGCCGCCGC | |
| **WZ0147_(podj)_reverse** | CCAATTGAGATCTGCTTAGCGCGCGTAGACCGACAGGC | |
| **WZ0148_(pcdf-mcherry)_forward** | GCCAATGGCGGCTACTAAGCAGATCTCAATTGGATATCGGCCGGC | pWZ064/ podj12 |
| **WZ0149_(pcdf-mcherry)_reverse** | GGTGTTCAGCAGCAGGGTGGCCGACCGGTGCTTGTA | |
| **WZ0150_(podj)_forward** | CACCGGTCGGCCACCCTGCTGCTGAACACCGACGACGG | |
| **WZ0151_(podj)_reverse** | CAATTGAGATCTGCTTAGTAGCCGCCATTGGCGGCGC | |
| **WZ0001_(pCDF-yfp)_forward** | CCTGTCGGTCTACGCGCGCCACCGGTCGGCCACCATGGT | pWZ065/ podj13 |
| **WZ0152_(pcdf-mcherryc)_reverse** | CCGTCGTCAGCGCGACCATTATATCTCCTTATTAAAGTTAAACAAAATTATTTCT | |
| **WZ0153_(podj)_forward** | CTTTAATAAGGAGATATAATGGTCGCGCTGACGACGGG | |
| **WZ0114_(pWZ013)_reverse** | GGTGGCCGACCGGTGGCGCGCGTAGACCGACAGGC | |
| **WZ0131_(yfp)_forward** | CAGCAGCCATCACCATCATCACCACAGCCAATGGTGAGCAAGGGCGAGGAGC | pWZ058 |
| **WZ0132_(yfp)_reverse** | CGTTTCGAATTCCATGGTGGCCGACCGGTGCTTGTACAGC | |
| **WZ0133_(divj)_forward** | CACCGGTCGGCCACCATGGAATTCGAAACGCTTCCAGACCCGT | |
| **WZ0134_(divj)_reverse** | CTTTCTGTTCGACTTAAGCATTATGCGGCCGCTCAGCGCGGCGCAAAGGCGA | |
| **WZ0154_(tipn-mchy)_forward** | CTTAGTATATTAGTTAAGTATAAGAAGGAGATATAATGAAGCCTAAGAAGCGCCAACC G | pWZ066 |
| **WZ0015_(hrsat-yfp)_reverse** | GTGGCCGGCCGATATCCAATTGAGATCTGCTTACTTGTACAGCTCGTCCATGCCG | |
| **WZ0025_(pxyfpc-6)_forward** | CTTAGTATATTAGTTAAGTATAAGAAGGAGATATAATGGTGAGCAAGGGCGAGGAGC | pWZ067 |
| **WZ0155_(cfp)_reverse** | CCACGACGGACTCCATGGTGGCCGACCGGTGCTTGTACAGC | |
| **WZ0156_(parb)_forward** | CACCGGTCGGCCACCATGGAGTCCGTCGTGGTGGG | |
| **WZ0157_(parb)_reverse** | GTGGCCGGCCGATATCCAATTGAGATCTGCTCAGATCCCGCGCGTCAGTCG | |
| **WZ0158_(ftsz)_forward** | CTTAGTATATTAGTTAAGTATAAGAAGGAGATATAATGGCTATTTCTCTTTCCGCGCCGC | pWZ068 |
| **WZ0159_(ftsz)_reverse** | GGTGGCCGACCGGTGGTTGGCCAGGCGGCGCAGGAACG | |
| **WZ0160_(mcherry)_forward** | CGCCGCCTGGCCAACCACCGGTCGGCCACCATGGTGAGC | |
| **WZ0015_(hrsat-yfp)_reverse** | GTGGCCGGCCGATATCCAATTGAGATCTGCTTACTTGTACAGCTCGTCCATGCCG | |
| **WZ0027_(pxchyc-6)_forward** | CTTAGTATATTAGTTAAGTATAAGAAGGAGATATAATGGTGAGCAAGGGCGAGGAGG | pWZ070 |
| **WZ0164_(mcherry)_(hrsat)_revers e** | CGAAAAGGGAAGAGAACATGGTGGCCGACCGGTGCTTGTACAGCTCGTCCATGCCGCC | |
| **WZ0165_(mreb)_forward** | CACCGGTCGGCCACCATGTTCTCTTCCCTTTTCGGCG | |
| **WZ0163_(mreb)_reverse** | GTGGCCGGCCGATATCCAATTGAGATCTGCCTAGGCCAGCGTGGATTCCAGG | |
| **WZ0170_(pbad-podj)_forward** | GGACGAGCTGTACAAGCACCGGTCGGCCACCATGAC | pWZ074 |
| **WZ0171_(pbad-podj)_reverse** | GCCCTTGCTCACCATTTAATTCCTCCTGTTAGCCCAAAA | |
| **WZ0172_(cfp)_forward** | GCTAACAGGAGGAATTAAATGGTGAGCAAGGGCGAGGAGC | |

| | | |
|---|---|---|
| **WZ0010_(PCDF-YFP)_reverse** | GGTGGCCGACCGGTGCTTGTACAGCTCGTCCATGCCG | |
| **WZ0173_(pbad)_forward** | CGAGCTGTACAAGTAAGAAGCTTGGCTGTTTTGGCGG | pWZ076 |
| **WZ0174_(pbad)_reverse** | CCCGTGTCTGCCCATTTAATTCCTCCTGTTAGCCCAAAAA | |
| **WZ0175_(plec-mcherry)_forward** | CTAACAGGAGGAATTAAATGGGCAGACACGGGGGGCC | |
| **WZ0176_(plec-mcherry)_reverse** | CCAAAACAGCCAAGCTTCTTACTTGTACAGCTCGTCCATGCCG | |
| **WZ282-PODJ14-F** | TGGCGGCTACTAAGCAGATCTCAATTGGAT | podj14 |
| **WZ283-PODJ14-R** | GATCTGCTTAGTAGCCGCCATTGGCGGCGC | |
| **WZ284-PODJ15-F** | CTTGGCCTTCTAAGCAGATCTCAATTGGAT | podj15 |
| **WZ285-PODJ15-R** | GATCTGCTTAGAAGGCCAAGGCCGAGCGGT | |
| **WZ286-PODJ18-F** | GCGCGCCGCCTAAGCAGATCTCAATTGGAT | podj18 |
| **WZ287-PODJ18-R** | GATCTGCTTAGGCGGCGCGCGGCGCGCCGG | |
| **WZ0184_(pcdf-mcherry)_forward** | GCGCCGCGCGCCGCCTAAGCAGATCTCAATTGGATATCGGCCGGC | PWZ079 /podj16 |
| **WZ0136_(pcdf-mcherry)_reverse** | CGGCGACGGGGTGGCCGACCGGTGCTTGTA | |
| **WZ0137_(podj08)_forward** | TACAAGCACCGGTCGGCCACCCCGTCGCCGCCGCCCGCGCAGG | |
| **WZ0185_(PODJ16)_reverse** | AATTGAGATCTGCTTAGGCGGCGCGCGGCGCGCCGG | |
| **WZ296-podj19-f** | GTCGGCCACCGAGCAGATCGCCGTCGCCGC | podj19 |
| **WZ297-podj19-r** | CGATCTGCTCGGTGGCCGACCGGTGCTTGT | |
| **WZ298-podj20-f** | GTCGGCCACCTTGCGCGCGCTTGAAGGCGC | podj20 |
| **WZ299-podj20-r** | GCGCGCGCAAGGTGGCCGACCGGTGCTTGT | |
| **WZ300-podj021-f** | GTCGGCCACCTTGGGCGCCGTCGAGACTGC | podj21 |
| **WZ301-podj021-r** | CGGCGCCCAAGGTGGCCGACCGGTGCTTGT | |
| **WZ302-podj022-f** | GTCGGCCACCAGCCAGGAACTGGTCGACCG | podj22 |
| **WZ303-podj022-r** | GTTCCTGGCTGGTGGCCGACCGGTGCTTGT | |
| **WZ304-podj023-f** | GGCGCGCTTGTAAGCAGATCTCAATTGGAT | podj23 |
| **WZ305-podj023-r** | GATCTGCTTACAAGCGCGCCTTCGACTTCT | |
| **WZ0193_(pbad)_forward** | ACGAGCTGTACAAGTAAGAAGCTTGGCTGTTTTGGCGG | pWZ082 |
| **WZ0194_(pbad)_reverse** | CCTGATGACGCGGTTTCATTTAATTCCTCCTGTTAGCCCAAAAA | |
| **WZ0195_(spmx-1-350)_forward** | GCTAACAGGAGGAATTAAATGAAACCGCGTCATCAGGTCTCCC | |
| **WZ0196_(spmx-1-350)_reverse** | GTGGCCGACCGGTGGTCCATCACGCCGACGGTTGC | |
| **WZ0197_(mcherry)_forward** | CGTCGGCGTGATGGACCACCGGTCGGCCACCATGGTGAGC | |
| **WZ0176_(plec-mcherry)_reverse** | CCAAAACAGCCAAGCTTCTTACTTGTACAGCTCGTCCATGCCG | |
| **WZ0193_(pbad)_forward** | ACGAGCTGTACAAGTAAGAAGCTTGGCTGTTTTGGCGG | pWZ083 |
| **WZ0198_(pbad-plec)_reverse** | GTGGCCGACCGGTGGGCCGCCACGAAGTCGCGAG | |
| **WZ0199_(yfp)_forward** | CGACTTCGTGGCGGCCCACCGGTCGGCCACCATGGTG | |
| **WZ0176_(plec-mcherry)_reverse** | CCAAAACAGCCAAGCTTCTTACTTGTACAGCTCGTCCATGCCG | |
| **WZ0173_(pbad)_forward** | CGAGCTGTACAAGTAAGAAGCTTGGCTGTTTTGGCGG | pWZ085 |
| **WZ0200_(pbad)_reverse** | GGATAAATCAAAGTTACGCATTTAATTCCTCCTGTTAGCCCAAAAA | |
| **WZ0201_(ibpa-yfp)_forward** | GGGCTAACAGGAGGAATTAAATGCGTAACTTTGATTTATCCCCGC | |
| **WZ0176_(plec-mcherry)_reverse** | CCAAAACAGCCAAGCTTCTTACTTGTACAGCTCGTCCATGCCG | |
| **WZ338-podj17-f** | AGCTGCGCCGCTCGTTGTCTTCGCCGCCGC | podj17 |
| **WZ339-podj17-r** | AGACAACGAGCGGCGCAGCTTCCAGCTTCC | |
| **WZ0219_(pcdf-mcherry)_forward** | CCGCGCGCCGCCTAATAAGCAGATCTCAATTGGATATCGGCCGGC | pWZ092/ podj24 |
| **WZ0220_(pcdf-mcherry)_reverse** | CGATGATCATTCGGTTGGTGGCCGACCGGTGCTTGTA | |
| **WZ0221_(podj36-702)_forward** | CACCGGTCGGCCACCAACCGAATGATCATCGAAGGCGATGG | |
| **WZ0222_(podj36-702)_reverse** | CAATTGAGATCTGCTTATTAGGCGGCGCGCGGCGCGC | |

| | | |
|---|---|---|
| **WZ0226_(pcdf-mcherry)_forward** | CCACGAACGTTCCAGCTAAGCAGATCTCAATTGGATATCGGCCGGC | pWZ094/ podj28 |
| **WZ0227_(pcdf-mcherry)_reverse** | GCAGTCTCGACGGCGCCCAAGGTGGCCGACCGGTGCTTGTA | |
| **WZ0228_(podj250-430)_forward** | CACCGGTCGGCCACCTTGGGCGCCGTCGAGACTGC | |
| **WZ0229_(podj250-430)_reverse** | CCAATTGAGATCTGCTTAGCTGGAACGTTCGTGGCGTTGG | |
| **WZ0230_(pcdf-mcherry)_forward** | GTCGAAGGCGCGCTTGTAAGCAGATCTCAATTGGATATCGGCCGGC | pWZ095/ podj25 |
| **WZ0231_(pcdf-mcherry)_reverse** | GATGATCATTCGGTTGGTGGCCGACCGGTGCTTGTA | |
| **WZ0221_(podj36-702)_forward** | CACCGGTCGGCCACCAACCGAATGATCATCGAAGGCGATGG | |
| **WZ0232_(podj36-635)_reverse** | CAATTGAGATCTGCTTACAAGCGCGCCTTCGACTTCTTG | |
| **WZ0235_(mcherry)_forward** | GAAATAATTTTGTTTAACTTTAATAAGGAGATATAATGGTGAGCAAGGGCGAGGAGG | pWZ099 |
| **WZ0236_(mcherry)_(hrsat)_revers e** | CGTTGTCGGTCGGCCGCATGGTGGCCGACCGGTGCTTGTACAGCTCGTCCATGCCGCC | |
| **WZ0237_(cpae)_forward** | CACCGGTCGGCCACCATGCGGCCGACCGACAACGA | |
| **WZ0238_(cpae)_reverse** | GGCTGTGGTGATGATGGTGATGGCTGCTGCCTACTTCTTCTTGAACAGGCCCGAGAACA TCG | |
| **WZ0239_(pled)_forward** | GAAATAATTTTGTTTAACTTTAATAAGGAGATATAATGAGCGCCCGGATCCTCGTCG | pWZ100 |
| **WZ0240_(pled)_reverse** | GGTGGCCGACCGGTGGGCGGCCTTGCCGACCACCG | |
| **WZ0241_(mcherry)_forward** | GTCGGCAAGGCCGCCCACCGGTCGGCCACCATGGTGAGC | |
| **WZ0242_(mcherry)_reverse** | GGCTGTGGTGATGATGGTGATGGCTGCTGCTTACTTGTACAGCTCGTCCATGCCGC | |
| **WZ0243_(popa)_forward** | GAAATAATTTTGTTTAACTTTAATAAGGAGATATAATGGCGGTTGACGCCCGAATCC | pWZ101 |
| **WZ0244_(popa)_reverse** | GGTGGCCGACCGGTGGCCCGCCTCGCGCTTCAGCG | |
| **WZ0245_(mcherry)_forward** | AAGCGCGAGGCGGGCCACCGGTCGGCCACCATGGTGAGC | |
| **WZ0242_(mcherry)_reverse** | GGCTGTGGTGATGATGGTGATGGCTGCTGCTTACTTGTACAGCTCGTCCATGCCGC | |
| **WZH452-podj29-f** | GGCCGGCGTGTAAGCAGATCTCAATTGGAT | podj29 |
| **WZH453-podj29-r** | GATCTGCTTACACGCCGGCCCCGAGCGCGC | |
| **WZH454-podj30-f** | GTCGGCCACCGGCGGCCTGCTGCTGCTGAA | podj30 |
| **WZH455-podj30-r** | GCAGGCCGCCGGTGGCCGACCGGTGCTTGT | |
| **WZ0294_(pWZ081)_forward** | TTTGATCGCGACGAACTCGTTGTCTTCGCCGCCGC | podj32 |
| **WZ0295_(pWZ081)_reverse** | GCCGGCGCGTCATCCGGCGCAGCTTCCAGCTTCC | |
| **WZ0296_(ped)_forward** | GCTGGAAGCTGCGCCGGATGACGCGCCGGCGGAGCC | |
| **WZ0297_(ped)_reverse** | GGCGAAGACAACGAGTTCGTCGCGATCAAACACCGGAGC | |
| **WZ0173_(pbad)_forward** | CGAGCTGTACAAGTAAGAAGCTTGGCTGTTTTGGCGG | pWZ121 |
| **WZ0298_(pbad)_reverse** | GGACCTTCTTCGTCATTTAATTCCTCCTGTTAGCCCAAAAA | |
| **WZ0299_(divk-mch)_forward** | GGGCTAACAGGAGGAATTAAATGACGAAGAAGGTCCTCATCGTGG | |
| **WZ0176_(plec-mcherry)_reverse** | CCAAAACAGCCAAGCTTCTTACTTGTACAGCTCGTCCATGCCG | |
| **WZ0173_(pbad)_forward** | CGAGCTGTACAAGTAAGAAGCTTGGCTGTTTTGGCGG | pWZ122 |
| **WZ0300_(pbad)_reverse** | GCAAGTCGGCCATTTAATTCCTCCTGTTAGCCCAAAAA | |
| **WZ0301_(ccka-mch)_forward** | CTAACAGGAGGAATTAAATGGCCGACTTGCAGCTCCAGG | |
| **WZ0176_(plec-mcherry)_reverse** | CCAAAACAGCCAAGCTTCTTACTTGTACAGCTCGTCCATGCCG | |
| **WZ0302_(pbad-mch)_forward** | GCCTTTGCGCCGCGCCACCGGTCGGCCACCATGGT | pWZ123 |
| **WZ0303_(pbad-mch)_reverse** | GGAAGCGTTTCGAATTCCATTTAATTCCTCCTGTTAGCCCAAAA | |
| **WZ0304_(divj)_forward** | GGGCTAACAGGAGGAATTAAATGGAATTCGAAACGCTTCCAGACCCGT | |
| **WZ0129_(divj)_reverse** | GGTGGCCGACCGGTGGCGCGGCGCAAAGGCGATGA | |
| **WZ0305_(pbad-mch)_forward** | CCCGAACTCGGCTTCCACCGGTCGGCCACCATGGT | pWZ124 |
| **WZ0306_(pbad-mch)_reverse** | GGTCGTACGAAGTCATTTAATTCCTCCTGTTAGCCCAAAA | |
| **WZ0307_(divl)_forward** | GGGCTAACAGGAGGAATTAAATGACTTCGTACGACCTGATCCTCGCG | |
| **WZ0017_(divl-cc-26-769)_reverse** | GGTGGCCGACCGGTGGAAGCCGAGTTCGGGCTGCA | |
| **WZH479-podj33-f** | AGCTGCGCCGGCCGCCGCGCGCGCCGCCGC | podj33 |

| | | |
|---|---|---|
| **WZH480-podj33-r** | GCGCGGCGGCCGGCGCAGCTTCCAGCTTCC | |
| **WZH489-plec1-f** | GAATTAAATGCAACGCGAGGCCATGGCCCA | plec1 |
| **WZH490-plec1-r** | CCTCGCGTTGCATTTAATTCCTCCTGTTAG | |
| **WZH491-plec2-f** | GCATCGCTTGATGATCCAGAGCCGCAAGGC | plec2 |
| **WZH492-plec2-r** | TCTGGATCATCAAGCGATGCACGCCAAAGG | |
| **WZH493-plec3-f** | GCTGCTGCTGGCCATCAAGACCCAGGAAGA | plec3 |
| **WZH494-plec3-r** | TCTTGATGGCCAGCAGCAGCGCCAGGGCGA | |
| **WZH495-plec4-f** | CGACATCACGCACCGGTCGGCCACCATGGT | plec4 |
| **WZH496-plec4-r** | CCGACCGGTGCGTGATGTCGGCGGCGGTCA | |
| **WZH501-podj31-f** | GAGCAGGCCCGCCTCGTTGTCTTCGCCGCCGCCGGCG | podj31 |
| **WZH502-podj31-r** | GAAGACAACGAGGCGGGCCTGCTCGATGATATCGCGG | |
| **WZ0308_(pbad-mch)_forward** | GCAGCTGCAGGCGGCGCACCGGTCGGCCACCATGGT | pWZ125 |
| **WZ0309_(pbad-mch)_reverse** | GCTGCAAGTCGGCCATTTAATTCCTCCTGTTAGCCCAAAA | |
| **WZ0310_(ccka-mch)_forward** | GCTAACAGGAGGAATTAAATGGCCGACTTGCAGCTCCAGG | |
| **WZ0203_(ccka)_reverse** | GGTGGCCGACCGGTGCGCCGCCTGCAGCTGCTGCTTGACG | |
| **WZ0173_(pbad)_forward** | CGAGCTGTACAAGTAAGAAGCTTGGCTGTTTTGGCGG | pWZ127 |
| **WZ0312_(pbad)_reverse** | GCGGCTCTGGATCATTTAATTCCTCCTGTTAGCCCAAAAA | |
| **WZ0315_(plec-pasc)_forward** | CTAACAGGAGGAATTAAATGATCCAGAGCCGCAAGGCCG | |
| **WZ0316_(plec-pasc)_reverse** | GGTGGCCGACCGGTGGGTGACGTCCAGCGCCACGC | |
| **WZ0317_(mcherry)_forward** | GCGCTGGACGTCACCCACCGGTCGGCCACCATGGTGAGC | |
| **WZ0176_(plec-mcherry)_reverse** | CCAAAACAGCCAAGCTTCTTACTTGTACAGCTCGTCCATGCCG | |
| **WZH521-plec-detald-f** | GGACGTCACCGCCATCAAGACCCAGGAAGA | pBAD-plec-ΔpasC |
| **WZH522-plec-detald-r** | TCTTGATGGCGGTGACGTCCAGCGCCACGC | |
| **WZH524-plec-detalc-f** | GAGCCGCAAGGAGGAGCGGATCGCCCAGGC | pBAD-plec-ΔpasC |
| **WZH525-plec-detalc-r** | TCCGCTCCTCCTTGCGGCTCTGGATCATCAG | |
| **WZ0173_(pbad)_forward** | CGAGCTGTACAAGTAAGAAGCTTGGCTGTTTTGGCGG | pWZ135 |
| **WZ0322_(pbad)_reverse** | CGATCCGCTCCTCCATTTAATTCCTCCTGTTAGCCCAAAAA | |
| **WZ0323_(plec-pasd-atg)_forward** | GCTAACAGGAGGAATTAAATGGAGGAGCGGATCGCCCAGG | |
| **WZ0320_(plec-pasd)_reverse** | GGTGGCCGACCGGTGCGTGATGTCGGCGGCGGTCATGACAAGACC | |
| **WZ0321_(mcherry)_forward** | GCCGCCGACATCACGCACCGGTCGGCCACCATGGTGAGC | |
| **WZ0176_(plec-mcherry)_reverse** | CCAAAACAGCCAAGCTTCTTACTTGTACAGCTCGTCCATGCCG | |
| **WZ0173_(pbad)_forward** | CGAGCTGTACAAGTAAGAAGCTTGGCTGTTTTGGCGG | pWZ136 |
| **WZ0312_(pbad)_reverse** | GCGGCTCTGGATCATTTAATTCCTCCTGTTAGCCCAAAAA | |
| **WZ0315_(plec-pasc)_forward** | CTAACAGGAGGAATTAAATGATCCAGAGCCGCAAGGCCG | |
| **WZ0320_(plec-pasd)_reverse** | GGTGGCCGACCGGTGCGTGATGTCGGCGGCGGTCATGACAAGACC | |
| **WZ0321_(mcherry)_forward** | GCCGCCGACATCACGCACCGGTCGGCCACCATGGTGAGC | |
| **WZ0176_(plec-mcherry)_reverse** | CCAAAACAGCCAAGCTTCTTACTTGTACAGCTCGTCCATGCCG | |
| **WZH526-plec-pasc-e-f** | CTAGCTAGCATGATCCAGAGCCGCAAGGC | pWZ129 |
| **WZH527-plec-pasc-e-r** | CGGGATCCTTAGGTGACGTCCAGCGCCACGC | |
| **WZH528-plec-pasd-e-f** | CTAGCTAGCGAGGAGCGGATCGCCCAGGC | pWZ130 |
| **WZH529-plec-pasd-e-r** | CGGGATCCTTACGTGATGTCGGCGGCGGTCA | |
| **WZH346-PTEV5-PODJ1-635-F** | GGCGCTAGCATGACGGCGGCTTCGCCATG | pwz91 |
| **WZH347-PTEV5-PODJ1-635-R** | GCGGATCCTTACAAGCGCGCCTTCGACTTC | |
| **WZH348-PTEV5-PODJ471-635-F** | GGCGCTAGCCCGTCGCCGCCGCCCGCGCA | pwz96 |

| | | |
|---|---|---|
| **WZH349-PTEV5-PODJ471-635-R** | CGCGGATCCTTACAAGCGCGCCTTCGACT | |
| **WZH352-PTEV5-PODJ250-430-F** | GGCGCTAGCTTGGGCGCCGTCGAGACTGC | pwz98 |
| **WZH353-PTEV5-PODJ250-430-R** | CGCGGATCCTTAGCTGGAACGTTCGTGGC | |
| **WZH497-pwz096-CYS-F** | TTTTCAGGGCTGTGCTAGCCCGTCGCCGCCGC | pwz96-cys |
| **WZH498-pwz096-CYS-R** | GGCTAGCACAGCCCTGAAAATACAGGTTTTC | |
| **WZH499-pwz098-CYS-F** | TTCAGGGCTGTGCTAGCTTGGGCGCCGTCGA | pwz98-cys |
| **WZH500-pwz098-CYS-R** | CAAGCTAGCACAGCCCTGAAAATACAGGTTTTC | |
| **WZH584-PopZ5-detalPED-F** | CATCTCGGAGGTCGCCGAGCAGCTGGTCGG | popz5 |
| **WZH585-PopZ5-detalPED-R** | GCTCGGCGACCTCCGAGATGATGCGTCGAA | |
| **WZH586-PopZ1-detalSA-F** | GGACGGTCGGTAAGAATTCGAGCTCGGCGC | popz1 |
| **WZH587-PopZ1-detalSA-R** | CGAATTCTTACCGACCGTCCTTGGGCATCA | |
| **WZ0368_(pcdf-mch)_forward** | CCCAAGGACGGTCGGTAAGAATTCGAGCTCGGCGC | popz3 |
| **WZ0369_(pcdf-mch)_reverse** | CGCCGGCGCGTCATCGGTGGCCGACCGGTGCTTGTA | |
| **WZ0370_(PopZ-PED-H2)_forward** | CACCGGTCGGCCACCGATGACGCGCCGGCGGAGCC | |
| **WZ0371_(PopZ-PED-H2)_reverse** | GAGCTCGAATTCTTACCGACCGTCCTTGGGCATCAGC | |
| **WZ0368_(pcdf-mch)_forward** | CCCAAGGACGGTCGGTAAGAATTCGAGCTCGGCGC | popz4 |
| **WZ0375_(pcdf-mch)_reverse** | CCAGCTGCTCGGCGACCTCCGAGATGATGCGTCGAATGG | |
| **WZ0373_(PopZ-H2)_forward** | CGCATCATCTCGGAGGTCGCCGAGCAGCTGGTCGG | |
| **WZ0374_(PopZ-H2)_reverse** | CGAGCTCGAATTCTTACCGACCGTCCTTGGGCATCAGC | |
| **podj-detalCC3-F** | GGATCAGCGCCAGGAACTGGTCGACCGCAT | PodJ-detal250-430 |
| **podj-detalCC3-R** | CCAGTTCCTGGCGCTGATCCAGGGCCTGGA | |
| **WZH624-PopZ2-detalH2-SA-F** | TCGCGACGAATAAGAATTCGAGCTCGGCGC | popz2 |
| **WZH625-PopZ2-detalH2-SA-R** | CGAATTCTTATTCGTCGCGATCAAACACCG | |
| **WZH628-POPZ6-F** | TCGCGACGAAACGCTGGAAGACGTCGTACG | popz6 |
| **WZH629-POPZ6-R** | CTTCCAGCGTTTCGTCGCGATCAAACACCG | |
| **WZH630-POPZ7-F** | GTCGGCCACCGATGACGCGCCGGCGGAGCC | popz7 |
| **WZH631-POPZ7-R** | GCGCGTCATCGGTGGCCGACCGGTGCTTGT | |
| **WZ0399_(pwz014)_forward** | GGTGTTTGATCGCGACGAAACGCTGGAAGACGTCGTACGCG | pWZ158/ popz8 |
| **WZ0369_(pcdf-mch)_reverse** | CGCCGGCGCGTCATCGGTGGCCGACCGGTGCTTGTA | |
| **WZ0370_(PopZ-PED-H2)_forward** | CACCGGTCGGCCACCGATGACGCGCCGGCGGAGCC | |
| **WZ0400_(ped)_reverse** | GACGTCTTCCAGCGTTTCGTCGCGATCAAACACCGGAGC | |
| **WZ0401_(pwz014)_forward** | GGTGTTTGATCGCGACGAATAAGAATTCGAGCTCGGCGC | pWZ159/ popz9 |
| **WZ0402_(pwz014)_reverse** | GCCGGCGCGTCATCGGTGGCCGACCGGTGCTTGTA | |
| **WZ0403_(ped)_forward** | GCACCGGTCGGCCACCGATGACGCGCCGGCGGAGCC | |
| **WZ0404_(ped)_reverse** | CCGAGCTCGAATTCTTATTCGTCGCGATCAAACACCGGAGC | |
| **WZ0113_(pwz013)_forward** | GGACGAGCTGTACAAGTAAGCAGATCTCAATTGGATATCGGCCGGC | pWZ161 |
| **WZ0405_(pacyc-tipn)_reverse** | GCCCTTGCTCACCATGGTGGCCGACCGGTGGGCCAGATCG | |
| **WZ0406_(ecfp)_forward** | CACCGGTCGGCCACCATGGTGAGCAAGGGCGAGGAGC | |
| **WZ0407_(ecfp)_reverse** | CCAATTGAGATCTGCTTACTTGTACAGCTCGTCCATGCCG | |
| **WZH678-pcdf-cfp-spmx(1-356)-f** | GGTGGAGCCGTAGGCAGATCTCAATTGGAT | pWZ201 |
| **WZH679-pcdf-cfp-spmx(1-356)-r** | GATCTGCCTACGGCTCCACCAGCGGCACGT | |
| **WZ0436_(pwz078)_forward** | GAGCGACGAAGAGTAGGCGGATTTGAACGTTGCGAA | pWZ170 |
| **WZ0171_(pbad-podj)_reverse** | GCCCTTGCTCACCATTTAATTCCTCCTGTTAGCCCAAAA | |

| | | |
|---|---|---|
| **WZ0434_(mch-podj)_forward** | AACAGGAGGAATTAAATGGTGAGCAAGGGCGAGGAGG | |
| **WZ0437_(pwz037)_reverse** | AACGTTCAAATCCGCCTACTCTTCGTCGCTCACATCGGGG | |
| **WZH684-podj42-702-f** | GTCGGCCACCGGCGATGGTCAGACCGCTGA | pWZ202 |
| **WZH685-podj42-702-r** | GACCATCGCCGGTGGCCGACCGGTGCTTGT | |
| **WZH686-podj1-601-f** | AGGCAAGGGCTAAGCAGATCTCAATTGGAT | pWZ203 |
| **WZH687-podj1-601-r** | GATCTGCTTAGCCCTTGCCTTCCGAGGCGG | |
| **WZ0438_(pcdf)_forward** | GGACGAGCTGTACAAGTAGGCAGATCTCAATTGGATATCGGC | pWZ171 |
| **WZ0439_(pcdf)_reverse** | CCGTGTCTGCCCATCTGGCTGTGGTGATGATGGTGATGG | |
| **WZ0440_(plec)_forward** | CCATCATCACCACAGCCAGATGGGCAGACACGGGGGGCC | |
| **WZ0252_(plec)_reverse** | GCCCTTGCTCACCATGGTGGCCGACCGGTGGGCCG | |
| **WZ0406_(ecfp)_forward** | CACCGGTCGGCCACCATGGTGAGCAAGGGCGAGGAGC | |
| **WZ0441_(ecfp)_reverse** | CCAATTGAGATCTGCCTACTTGTACAGCTCGTCCATGCCG | |
| **WZH694-pcdf-cfp-spmx1-155-f** | CGGCGAATGGTAGGCAGATCTCAATTGGAT | pWZ204 |
| **WZH695-pcdf-cfp-spmx1-155-r** | GATCTGCCTACCATTCGCCGTTGGCCGGCG | |
| **WZ0442_(pcdf-cfp)_forward** | CCGCTGGTGGAGCCGTAGGCAGATCTCAATTGGATATCGGC | pWZ172 |
| **WZ0443_(pcdf-cfp)_reverse** | CGCGGGGACGGTGGCCGACCGGTGCTTGTA | |
| **WZ0444_(spmx156-356)_forward** | TACAAGCACCGGTCGGCCACCGTCCCCGCGCCCAGCCCCGT | |
| **WZ0445_(spmx156-356)_reverse** | AATTGAGATCTGCCTACGGCTCCACCAGCGGCACGT | |
| **WZ0050_(PBAD)_forward** | GGTCTACGCGCGCTAAGAAGCTTGGCTGTTTTGGCGG | pWZ173 |
| **WZ0186_(pcdf-mcherry)_reverse** | CGAAGCCGCCGTCATGGTGGCCGACCGGTGCTTGTA | |
| **WZ0187_(PODJ1-470)_forward** | CACCGGTCGGCCACCATGACGGCGGCTTCGCCATGG | |
| **WZ0446_(podjdeltaPSE)_reverse** | CAAAACAGCCAAGCTTCTTAGCGCGCGTAGACCGACAGG | |
| **WZ0447_(pcdf-cfp)_forward** | CCGCTGGTGGAGCCGCACCGGTCGGCCACCATGGTGAGC | pWZ174 |
| **WZ0448_(pcdf-cfp)_reverse** | CGCGGTTTCATCTGGCTGTGGTGATGATGGTGATGGC | |
| **WZ0449_(spmx1-356)_forward** | CCATCATCACCACAGCCAGATGAAACCGCGTCATCAGGTCTCCCG | |
| **WZ0450_(spmx1-356)_reverse** | GGTGGCCGACCGGTGCGGCTCCACCAGCGGCACGTCC | |


Bowman, G.R., Perez, A.M., Ptacin, J.L., Ighodaro, E., Folta-Stogniew, E., Comolli, L.R., and Shapiro, L. (2013). Oligomerization and higher-order assembly contribute to sub-cellular localization of a bacterial scaffold. Mol Microbiol *90*, 776-795.

Holmes, J.A., Follett, S.E., Wang, H., Meadows, C.P., Varga, K., and Bowman, G.R. (2016). Caulobacter PopZ forms an intrinsically disordered hub in organizing bacterial cell poles. Proc Natl Acad Sci U S A *113*, 12490-12495.

Huitema, E., Pritchard, S., Matteson, D., Radhakrishnan, S.K., and Viollier, P.H. (2006). Bacterial birth scar proteins mark future flagellum assembly site. Cell *124*, 1025-1037.

Ptacin, J.L., Gahlmann, A., Bowman, G.R., Perez, A.M., von Diezmann, A.R., Eckart, M.R., Moerner, W.E., and Shapiro, L. (2014). Bacterial scaffold directs pole-specific centromere segregation. Proc Natl Acad Sci U S A *111*, E2046-2055.

Radhakrishnan, S.K., Thanbichler, M., and Viollier, P.H. (2008). The dynamic interplay between a cell fate determinant and a lysozyme homolog drives the asymmetric division cycle of Caulobacter crescentus. Genes & development *22*, 212-225.

Thanbichler, M., Iniesta, A.A., and Shapiro, L. (2007). A comprehensive set of plasmids for vanillate- and xylose-inducible gene expression in Caulobacter crescentus. Nucleic Acids Research *35*.

Viollier, P.H., Sternheim, N., and Shapiro, L. (2002). Identification of a localization factor for the polar positioning of bacterial structural and regulatory proteins. Proc Natl Acad Sci U S A *99*, 13831-13836.

Wheeler, R.T., and Shapiro, L. (1999). Differential localization of two histidine kinases controlling bacterial cell differentiation. Molecular cell *4*, 683-694.
